# The retinal pigment epithelium undergoes anisotropic stretching and nuclear size scaling during optic cup morphogenesis in a fish model

**DOI:** 10.64898/2026.07.12.737769

**Authors:** François Agnès, Mathilde Pain, Dimitri Vérité, Parisa Zia, Estelle Giry, Jorge Torres-Paz, Sylvie Rétaux

## Abstract

The morphogenesis of the optic cup provides a robust system for studying how two apposed epithelial monolayers with distinct properties fold and stretch in a coordinated manner to form the primordial eye. While much research has been conducted on the temporal dynamics of retinal neuroepithelium invagination, the spatial organization and stretching of the retinal pigment epithelium has received less attention. The fish species *Astyanax mexicanus* offers a unique model to examine the mechanisms of optic tissue morphogenesis through a comparative lens, as it exhibits natural variation in eye development between its river-dwelling and cave-adapted morphs. Using quantitative 3D imaging of optic cups from both morphs, we found that RPE morphogenesis involves transient, graded, and anisotropic cell stretching that patterns the epithelium during optic cup shaping. Analyses of RPE nuclear spacing and cell morphology showed that tissue stretching gradually increases along the proximo-distal axis, suggesting maximal tension in the elongated distal RPE cells aligned along the optic cup meridians. Furthermore, nuclear volumes and apical surface areas of RPE cells scaled spatially along the same axis, independently of endoreplication. In the cavefish natural mutant, RPE expansion was delayed by over six hours and proximal stretching exhibited altered isotropy, indicative of disrupted temporal coordination and suggesting modified mechanical constraints. These results demonstrate that RPE morphogenesis is a highly heterogeneous process from a spatiotemporal perspective, offering new insights into the study of the biomechanical principles of eye development in vertebrates.

**Summary statement:** This study reveals the emergence of cell morphology gradients within the retinal pigment epithelium during morphogenesis of the eye in two distinct populations of the same species of fish.

## Introduction

Epithelial monolayers form sheets of polygonal cells that cover the external or internal surfaces of organs in animals. These sheets are composed of cells with diverse geometries (e.g. cuboidal, columnar, prismatic, squamous) (Lemke and Nelson, 2021; Gómez-Gálvez et al., 2021), organized according to planar cell polarity (Goodrich and Strutt, 2011; Wallingford, 2012) and polarized along an apico-basal axis (Buckley and St Johnston, 2022). During development, epithelial monolayers undergo profound shape changes while preserving cohesion and barrier integrity. These transformations rely on the dynamic remodeling of intercellular junctions, allowing cells to rearrange and deform without compromising tissue continuity (Mira-Osuna and Le Borgne, 2024). Forces are generated by molecular motors, notably actomyosin networks, and transmitted through cytoskeletal elements to cell-cell adhesive complexes (Munjal & Lecuit, 2014; Murrell, 2020). Epithelial cells exhibit elastic properties that allow them to deform in response to external forces or to induce the displacement of neighbouring cells (Bonnet et al., 2012; Bouzignac and Suzanne, 2025). Tissue mechanical forces and the resulting stresses (e.g. compression, stretching) contribute to determining cell shape and packing (Graner and Riveline, 2017). Tissue deformation is driven by the interplay between deterministic genetic programs and self-organized mechanical processes integrating biochemical, geometric, and physical information (Collinet and Lecuit, 2021). Tissue morphogenesis can thus be viewed as the outcome of dynamic changes in cell size, shape, number, position, and gene expression (Heisenberg and Bellaïche, 2013; Heer et al., 2017). Folding of epithelial monolayers involves a coupling between active bending and tissue tension, with mechanical asymmetries generated either locally or across the tissue, and requiring non-uniform mechanical loads or geometric constraints (Fouchard et al., 2020; Tozluoǧlu and Mao, 2020; Wang, 2021).

The morphogenesis of the vertebrate optic cup (OC) is a powerful system for investigating how two connected, facing epithelial monolayers with distinct organizations fold and stretch to shape the primordial eye. The optic primordium comprises a pseudo-stratified retinal neuroepithelium (RNE), from which all retinal neurons originate, and, in its continuity, a prospective retinal pigment epithelium (RPE), which acquires essential visual and trophic support functions (Fuhrmann, 2010) as well as tremendous regenerative potential (George et al., 2021). These two epithelial domains originate from the same anterior medial eyefield in the anterior neural plate, which evaginates bilaterally to form the optic vesicles (England et al., 2006; Rembold et al., 2006). Continued tissue elongation through extended evagination (Kwan et al., 2012) establishes a polarized bilayer optic vesicle in which the future RPE is specified in the dorso-posterior region and expands (Cechmanek and McFarlane, 2017). Subsequent movements and local cell shape changes transform this vesicle into a hemispheric OC composed of an inner retinal epithelium and an outer RPE layer (Casey et al., 2021; Norden, 2023). This morphogenetic sequence is broadly conserved across vertebrates (Cardozo et al., 2023), although its dynamics differs between rapidly developing teleosts and slower-developing amniotes (Moreno-Mármol et al., 2021).

Recent studies in zebrafish have revealed coordinated epithelial cell flow from the outer to the inner layer of the RNE OC around the ventricular ridge (rim movement). This process actively translocates a substantial number of cells into the RNE around much of the optic vesicle circumference (Kwan et al., 2012; Heermann et al., 2015, Sidhaye and Norden, 2017, Soans et al., 2022). Quantitative analyses of invagination mechanics have shown that basal constriction of lens-facing RNE cells drives tissue folding, while actomyosin oscillations generate mechanical forces transmitted via extracellular matrix attachments (Nicolás-Pérez et al., 2016; Sidhaye and Norden, 2017). Inhibition of basal actomyosin contraction delays, but does not prevent RNE invagination, suggesting the involvement of additional mechanisms (Sidhaye and Norden, 2017). Interestingly, studies in zebrafish indicate that RNE invagination can take place even under reduced cell proliferation, suggesting that other additional processes participate in OC folding (Sidhaye and Norden, 2017).

Studies in teleosts have revealed that RPE expansion to cover the back of the entire prospective retina occurs in two phases: an initial increase in cell number through progenitor specification, largely independent of proliferation, followed by cell flattening and stretching along the proximo-distal axis accompanying OC invagination (Kwan et al., 2012; Cechmanek and McFarlane, 2017; Moreno-Mármol et al., 2021). Local and specific perturbations of the RPE cytoskeleton impair tissue stretching and compromise OC folding, suggesting that the RPE contributes actively to morphogenesis by generating tissue-autonomous deformation through intracellular cytoskeletal remodelling (Moreno-Mármol et al., 2021). These findings raise key questions: How do RPE cells deform in space and time? Is stretching uniform? What is the mechanical role of the RPE?

The Mexican tetra *Astyanax mexicanus*, with its surface-dwelling (SF) and cave-adapted (CF) morphs (Jeffery, 2020), provides a comparative framework for investigating RPE morphogenesis. Although eye morphogenesis in SF is similar to that in zebrafish (Devos et al., 2021), CF embryos exhibit a series of defects including smaller eyes and altered tissue geometry (Figure 1, schematic left). The optic vesicles elongate but remain smaller following evagination, and OC invagination is temporarily impaired, contributing to coloboma (Devos et al., 2021). While RPE identity is specified and maintained in CF during optic morphogenesis, its expansion to cover the whole RNE shows a delay compared to SF (Devos 2021). This natural variation offers an original paradigm for the dissection of the spatial and temporal coordination and the mechanics of RPE morphogenesis under evolutionary and developmental constraints.

**Figure 1:**
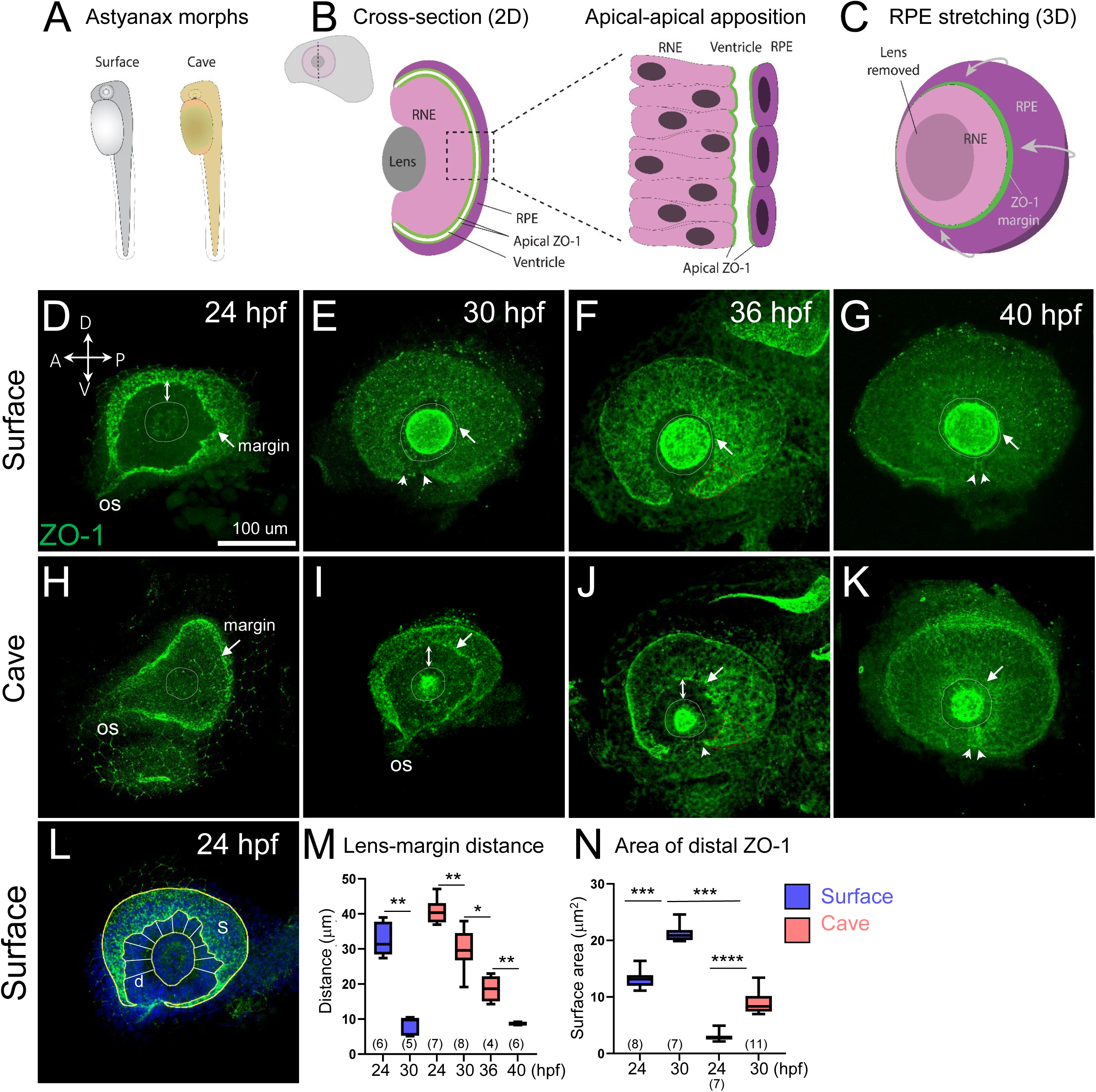
Spreading of the RPE is delayed in cavefish. (A) Embryos of the two morphs are represented. (B) The scheme (top left) shows a lateral view of the head and OC from which is a cross-section was drawn. The transversal cross section represents the organisation of the OC during folding. The two green lines indicate the apical side of the RPE and the RNE accumulating ZO-1. The white line indicate the ventricle. The drawing on the right represents the global morphologies of cells from the pseudostratified RNE and the flattened RPE, separated by a ventricle. (C) A 3D distal, slightly tilted view of the OC is shown, summarising data from Cechmanek et al., (2017) and Moreno-marmol et al., (2021). In the second phase of expansion, the RPE stretches around the prospective neural retina from the inner leaflet of the eye vesicle in all axes to abut the lens. (D-L) Whole-mount immunostaining for ZO-1 in dissected OC from SF and CF at the indicated developmental stages, shown in distal-up anterior left orientation (D). Confocal images correspond to maximum intensity projections. ZO-1 labels the apical junctions of both the RPE and the connected and facing RNE. The dashed circles delineates the lens contour. Arrows in magenta mark the distal edge of the ventricle, and double arrows indicate the distance between the distal ZO-1 margin and the lens border. Arrowheads indicate the limits of the optic fissure. Position of the optic stalk (OS) is indicated. Scale bar: 100 μm. (I) Schematic overlaid on a 24 hpf SF image illustrating the two quantitative parameters used to assess RPE expansion: the distance (d) from the distal ZO-1 margin to the lens, and the distal surface area (S) corresponding to the projected ZO-1-labelled region. Circular lines indicate that multiple measurements were taken to calculate the mean d value for each sample. (M,N) Quantification of d and S parameters in both morphs. The number of stars above each line indicates the corresponding *P*-value for the comparison.

Here, using quantitative 3D imaging, nuclear morphometric, and cell-shape analysis, we show that normal RPE morphogenesis involves anisotropic stretching along the proximo-distal and dorso-ventral axes of the OC, with maximal expansion in the proximal region and high density and cell elongation in the distal region. Stretching is temporally restricted to the period of OC shaping and resumes afterwards. Strikingly, we observed nuclear enlargement, which does not result from endoreplication. In the CF natural mutant, we observed a delay of over six hours in RPE expansion, and a disorganized proximal RPE showing higher relative stretching, compared to the SF. Our findings reveal previously unrecognized spatiotemporal coordination in RPE morphogenesis, highlighting the interplay between timing, tissue mechanics, and regional specialization in a conserved vertebrate morphogenetic program.

## Results

### RPE spreading is heterochronic in cavefish

To compare OC folding dynamics between the two *Astyanax* morphs, we stained whole-mount embryos between 24 and 40 hpf (hours post-fertilization) with an antibody against ZO-1, a tight junction protein that labels the apical cell membranes of epithelial cells (Zihni et al., 2016). To visualize RPE expansion, we dissected the OC and mounted them in lens-facing distal view (Figure 1, schematic). At these stages, ZO-1 labelled the apical membranes of both the RPE and the RNE, which are separated by a thin ventricle (Figures 1A-H, and Movie 1). The two epithelial layers facing each other were very close, as revealed by an average inter-apical space of slightly more than 2 µm in the SF that was twice larger in the CF (Fig. S1 and Figure 2DH). The accumulation of ZO-1 at the circumferential edge of the ventricle formed a distinct border enabling us to quantify the extent of RPE expansion, using two parameters: (i) the mean distance (d) between the distal margin of the ZO-1 staining and the lens border; and (ii) the area (S) covered by the distal RPE (Figure 1L). In SF at 24hpf, the ZO-1 margin was irregular and located on average 30 microns (d) away from the lens (Figure 1D and IJ) and the RPE already covered a substantial portion of the distal OC (OC; Figure 1N). At 30 hpf, the ZO-1 margin had reached the lens, with a 1.6-fold increase of the S parameter (Figure 1E and JK), indicating near completion of the RPE spreading in SF, seen as well in later stages (Figure 1FD). Compared to SF and as expected (Devos et al., 2021), the OC in CF showed a reduced size, a distinct shape, a larger optic stalk and an opened optic fissure phenotype (Figure 1H-G). At 24 hpf, ZO-1 labelling defined a spoon-like 3D shape in CF and the area S was smaller than in SF (Figure 1H, JK and Movie 2). This suggested that expansion of the RPE was significantly delayed in CF. At 30 hpf however, the ZO-1 labelling in CF resembled that of SF at 24 hpf (Figure 1D,F), with an irregular ZO-1 margin and a similar distance from the lens (d) (Figure 1I,J). Nevertheless, in CF the S parameter showed a 3-fold increase between 24 hpf and 30 hpf (Figure 1N), indicating the progression of RPE expansion. Interestingly, both the d and the S parameters showed higher variance in CF, indicating high inter-individual variability at 30 hpf. At 36 hpf, the ZO-1 margin was closer to the lens (Figure 1J,J) but the ventral edges of the ZO-1 labelling were not joined in CF (Figure 1J, arrowheads), typical of the coloboma phenotype in CF (Devos et al., 2021). Ventrally, the ZO-1 labelling delineated reduced size and different OC shape compared to SF at 36 hpf and to CF at 30 hpf (compare Figure 1J to Figure 1F and 1F, red dotted line). This suggested asymmetric and heterochronic RPE expansion in the ventro-distal region (perifissural) compared to the dorso-distal region. At 40 hpf, the distal margin of the RPE had finally reached the lens in CF (Figure 1K,J).

**Figure 2:**
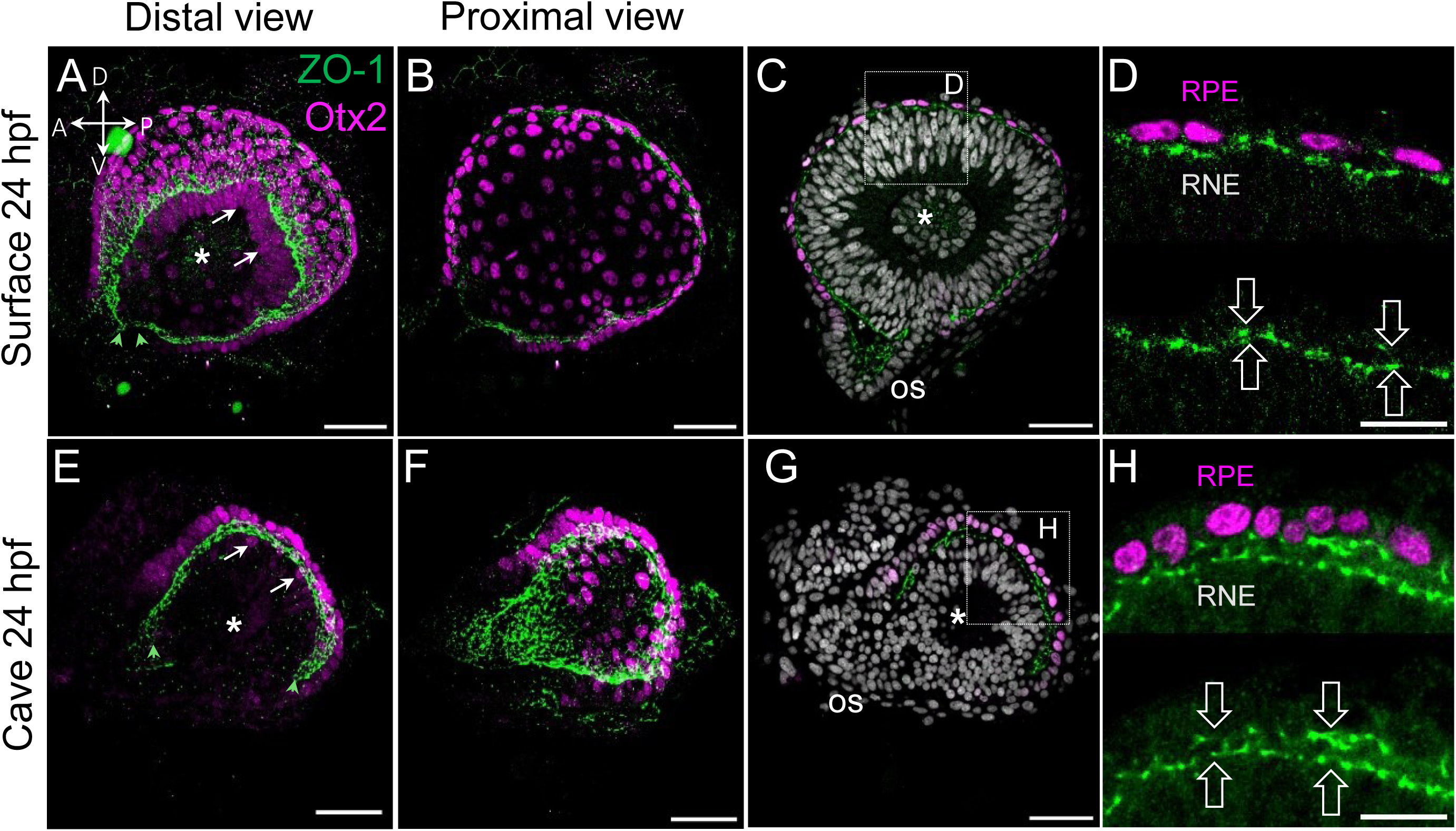
Comparative anatomy of the RPE in SF and CF at an equivalent developmental stage. (A–H) Confocal images of dissected SF and CF OC at 24hpf, immunostained for ZO-1 (green) and Otx2 (magenta). (A–B, E–F) 3D renditions (3D viewer) showing the organization of the RPE nuclei in the distal and proximal regions. Orientation of the OC in (A) applies to panels A-G. (A,E) The ZO-1 ventricle margin allows distinguishing the distal RPE nuclei from the uppermost distal RNE nuclei, likely corresponding to ciliary marginal zone progenitors (arrows). Arrowheads in indicate the position of the most ventral distal end of the ventricle. (C, D, G, H) Maximum intensity projections highlight the intermediate zone. ZO-1 staining outlines the circumferential ventricle, distinguishing outer RPE nuclei from inner facing RNE nuclei. (D,H) Zoom of insets shown in C and G, respectively. Symmetric wide arrows in (D,H) point to the apical surface of the two epithelia. The OS and asterisks, respectively, indicate the position of the optic stalk region and the centre of the lens. Scale bar: 20 µm.

In sum, outlining the position of the circumferential ventricular edge during OC folding with ZO-1 staining provides a quantitative readout of the second phase of RPE expansion and reveals a delay of more than six hours between the two *Astyanax* morphs. Because RPE flattening and OC folding occurs simultaneously and are linked through a shared ventricular geometry, comparable extents of RPE advancement reflect equivalent global states of OC folding. Consequently, the advancement of expansion of the RPE is similar in 24 hpf SF and 30 hpf CF, respectively (Figure 1L).

RPE anatomy during OC folding

Time-lapse analyses have described RPE expansion to occur simultaneously along several axes of the OC (Kwan et al., 2012; Cechmanek and McFarlane, 2017; Moreno-Mármol et al., 2021). To determine whether regional differences accompany RPE expansion, we examined its organization in fixed, dissected OCs. We immunostained embryos to detect ZO-1 together with Otx2, one of the key transcription factors regulating RPE specification and differentiation in vertebrates (Martinez-Morales et al., 2017; Bharti et al., 2012; Iwai-Takekoshi et al., 2016). We then compared different regions of the RPE in the two morphs at the same developmental stage (24 hpf) (Figure 2). At this stage, Otx2 labelled all RPE nuclei in the two morphs specifically (Movie 3 and Movies 4A and 4B). The circumferential edge of the ventricle delimited by ZO-1 labelling enabled us to distinguish RPE nuclei from those of adjacent RNE cells (Figure 2AD, arrows, Movies 3 and 4). This was particularly useful in SF, where the most distal, elongated RNE nuclei were also Otx2-positive (they likely corresponded to progenitor cells of the ciliary marginal zone Figure 2A, asterisk and Movie 3). In contrast, in CF, few and more rounded Otx2-positive nuclei were observed in the most distal region of the RNE, likely representing an earlier stage of differentiation (Figure 2E, asterisk).

Surprisingly, we observed regional differences in RPE nuclei distribution and morphology (Movie 3). In particular, the density of RPE nuclei seemed higher in the distal region close to the ZO-1 margin (Figure 2A,E and Fig. S2) than in the proximal region at the back of the prospective retina (Figure 2B,F), suggesting differential stretching within the RPE along its proximo-distal axis, in the two morphs. Furthermore, in the transition zone between the proximal and distal regions of the RPE, as shown in parasagittal optical sections, the nuclei in the SF were flattened (Figure 2CD), whereas those in the CF appeared more rounded (Figure 2GH).

To discard a potential contribution of cell proliferation to RPE expansion and proximo-distal heterogeneity in RPE nuclei distribution, we quantified mitotic activity using PH3 immunostaining combined with Otx2 to identify RPE cells. At 24 hpf, mitotic figures were scarce in both morphs, consistent with the low proliferative activity previously reported for the zebrafish RPE during expansion (Kwan et al., 2012; Cechmanek and McFarlane, 2017; Moreno-Mármol et al., 2021). Quantification of PH3⁺/Otx2⁺ nuclei revealed no significant difference between distal and proximal regions, indicating spatial homogeneity of proliferation within the RPE (data not shown).

These results indicate that the RPE proliferates at a low rate during OC shaping in the Mexican Tetra, and that regional differences in RPE expansion do not arise from differential cell division rates. To gain a better understanding of RPE stretching, we next quantified several parameters of RPE nuclear and cell organization and morphology at the whole-tissue and local scales.

### Nuclear organization varies along the proximo-distal axis during RPE spreading

Assuming that internuclear distance reflects RPE expansion due to stretching, we compared this parameter between the two morphs at the same embryonic stage (SF and CF at 24 hpf) and at similar stage of OC development (SF 24 hpf *vs* CF 30 hpf). To this end, we used Imaris software to create spots corresponding to the positions of RPE nuclei from the Otx2 channel (Figures 3A-C, Movie 5). We then used the x, y, z coordinates of these spots to calculate distance to 3 nearest neighbours (NND3) and modelled the spatial distribution of NND3 along the proximal-distal axis through colour-coding (Figure 3A’-C’). The quantification revealed a significant asymmetry in the distribution of NND3 for both morphs at each stage (Figure 3D, Movie 5). At 24 hpf in SF, the average nuclear NND3 in the proximal region was 1.7 times greater than that in the distal region (mean: 17.3 µm *vs* 10,2 µm), indicating more intense stretching in the proximal region (Figure 3D). At this stage in CF, the NND3 value was also higher in the proximal region than in the distal region (fold change ×1.4, mean: 12.5 µm *vs* 8.8 µm). There was also a slight, but significant difference in the distal region compared to SF (Figure 3D), which indicated a general reduction in stretching in the CF natural mutant. At 30 hpf in CF, the NND3 of the proximal nuclei was higher than that of the distal nuclei (×1.6). The mean proximal NND3 value was similar to SF at 24 hpf, but showed increased interindividual variability (Figure 3D). These results further indicated that RPE stretching begins in the proximal region, and strengthened the hypothesis that RPE stretching is delayed in CF. In CF, proximal NND3 increased by 20% between 24 and 30 hpf, compared to 7% in the distal region, also suggesting a gradual increase in RPE stretching along the distal-to-proximal axis.

**Figure 3.**
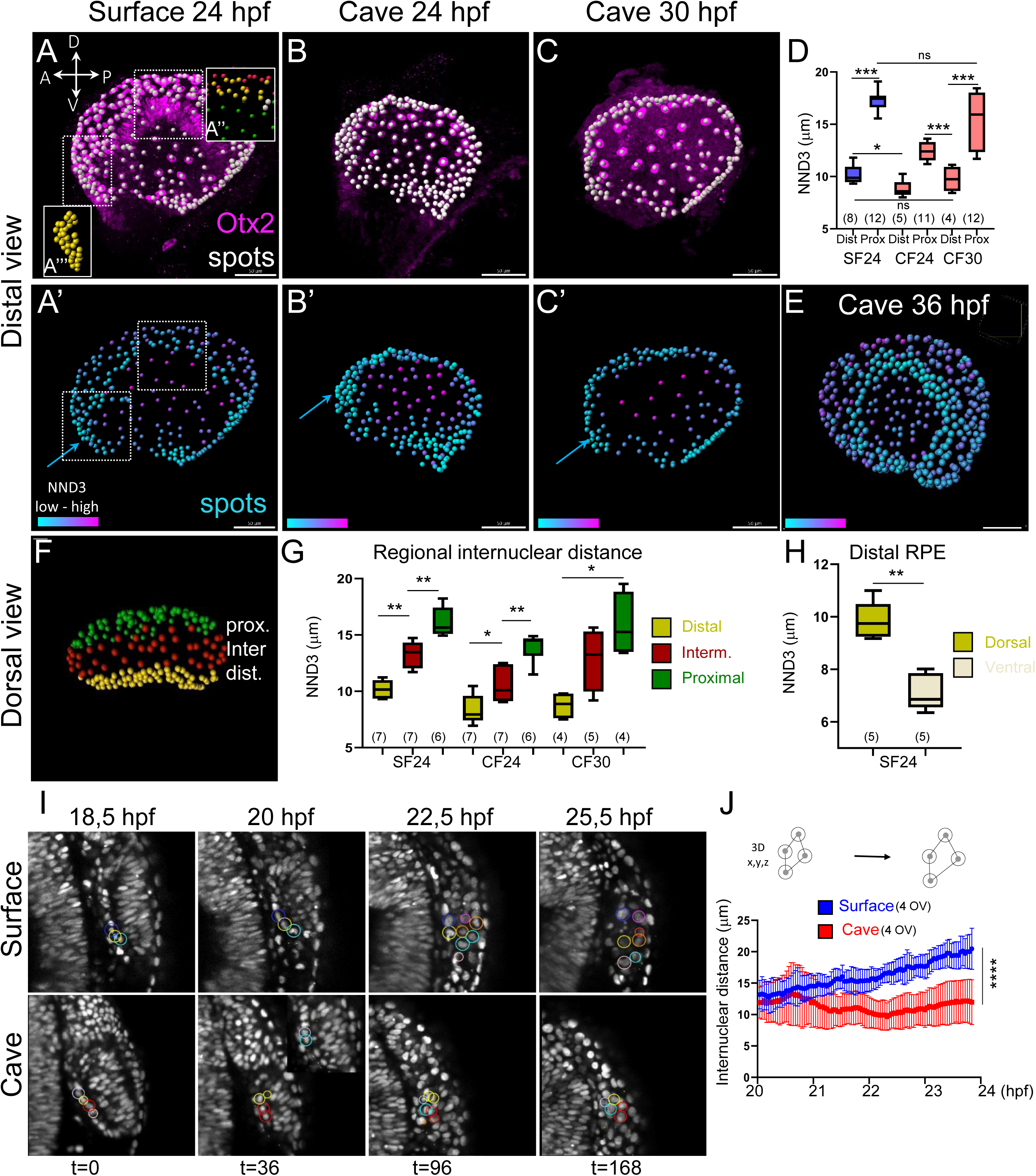
Heterogeneities in nuclear organization along the proximo-distal and dorso-ventral axes during RPE spreading. (A–C) 3D views of the RPE from dissected SF and CF OC at the indicated stages, immuno-stained for Otx2 (RPE nuclei, magenta). White dots correspond to the RPE nuclei only (Imaris spots). Orientation of the OC in (A) applies to panels A-C. The orientation shown in (A) applies to all subsequent panels. Scale bar 50 µm. (D) Comparison between distal and proximal RPE regions of the NND3 parameter at the indicated stages and morphs. (A’-C’,E) Colour-coded 3D maps of nearest-neighbour distances (NND3) across the RPE surface for the same sample as in (A-C) and CF at 36 hpf (E). The colour scale reveals graded patterns of nuclear density along the proximo-distal axis. (F) Colour-coded 3D map of NND3 across the RPE of surface fish in dorsal view highlighting the distal, intermediate and proximal regions. (G) Quantifications of NND3 across different RPE sub-regions shown in F. (H) Quantifications of NND3 across the dorsal and ventral regions of the distal RPE. The regions used for quantification (yellow dots) are shown in A (insets A’’ and A’’’). Coloured dots in A’’ distinguish the distal (yellow), intermediate (green) and proximal (red) regions. (I) Dorsal views of the same OC at the indicated developmental stages, shown at maximum intensity projection for both morphs. Circles highlight individual nuclei tracked during the developmental window. Red, orange and pink circles in SF lying that lie in other planes are not shown at t36 and t96. The apparent displacement of nuclei across different focal planes reflects their relative movement over time. The scale is identical in all cave and surface panels. (J) Quantification of internuclear distance between pairs of RPE nuclei over 5 hours. The graphic illustrates the vectors between the 4 adjacent RPE nuclei used for quantification and the progressive increase in length over time.

To better characterize organization of the nuclei along the proximo-distal axis, we also compared the NND3 in the proximal, intermediate/transition zone (where the OC curvature is pronounced) and distal regions (Figure3 F-G and Movie 6). This was done in the dorsal region of the OC to avoid bias due to the coloboma (failure of optic fissure closure) in the ventral OC in CF. To this end, we defined subsets of consecutive nuclei (Figure 3A, inset A’’ and Figure 1I). As shown by the quantifications, NND3 increased linearly along the distal-to-proximal axis in both morphs at each stage (Figure 3G and Fig. S3), indicating graded stretching along this axis.

Interestingly, while the RPE nuclei near the edge of the ventricle appeared highly compact in the SF at 24 hpf, we observed a difference in nuclear organisation between the ventral and dorsal parts of the distal RPE (Figure 3A: yellow spots in insets A’’ and A’’’; Figure 3A’: turquoise spots in square). As shown in Figure 3H, the ventral nuclei were more densely packed than the dorsal nuclei, suggesting a delay in the timing of tissue stretching in the region close to the optic fissure compared to dorsal regions.

Overall, these results demonstrate that stretching of the RPE occurs in a patterned manner along the proximal-distal axis (and the dorso-ventral axis to a lesser extent). Area expansion in the proximal region is the more pronounced, contrasting with the distal region that initially exhibits a compact arrangement of nuclei, suggesting that the two regions are subject to different mechanical constraints.

### RPE expansion dynamics: insights from live imaging

To compare dynamic changes in RPE spreading during OC folding between the two morphs, we re-analysed an existing live imaging dataset (Devos et al., 2021). Using the Mastodon plugin in Fiji (Girstmair et al. 2025), we tracked a subset of putative RPE nuclei and reconstructed their 3D trajectories over ∼7 hours of development between 18.5hpf and 25.5hpf, in both morphs (Figure 3I). Tracking of proximal RPE nuclei was not feasible due to insufficient spatial resolution in this region, thus we tracked distal RPE nuclei. As in the analyses of fixed tissue, we used variations in internuclear distances between several pairs of neighbouring RPE nuclei as an indicator of the expansion dynamics of the RPE. In SF, nuclei progressively dispersed over time, resulting in a gradual increase in internuclear spacing (Figure 3J). In contrast, RPE nuclei showed significantly reduced dispersion in CF. These nuclei, which migrated at the same speed than in SF (Fig. S4), remained more closely clustered throughout the same imaging period, with higher variability between pairs of neighbouring cells (Figure 3J).

These live imaging data indicate progressive RPE expansion in SF between 20 and 24 hpf, whereas expansion appears significantly reduced and temporally altered in CF during the same developmental window.

### Proximal RPE stretching is transient

Is RPE stretching temporally regulated? To examine this possibility, we compared the nuclear organization of the two morphs at 72 hpf, when the retina has become layered (Fig. S5). At this stage, the proximal RPE exhibited relatively compact and homogeneous nuclear organization in both morphs (Figure 4AB). In SF, the average NND3 value was 1,7 fold lower than at 24hpf (Figure 4C and Figure 3D). This indicated that proximal cells had not only ceased to stretch but also had reduced their stretching level by this advanced stage, and this was also observed in CF (data not shown). In addition, we observed that SF proximal RPE nuclei were arranged in a highly regular, ordered manner (Figure 4D). By contrast, this organization was often disrupted in CF, resulting in clustering and spacing (Figure 4E, magenta and yellow dotted line), whilst individual nuclear size was comparable in the two morphs (Figure 4F).

**Figure 4:**
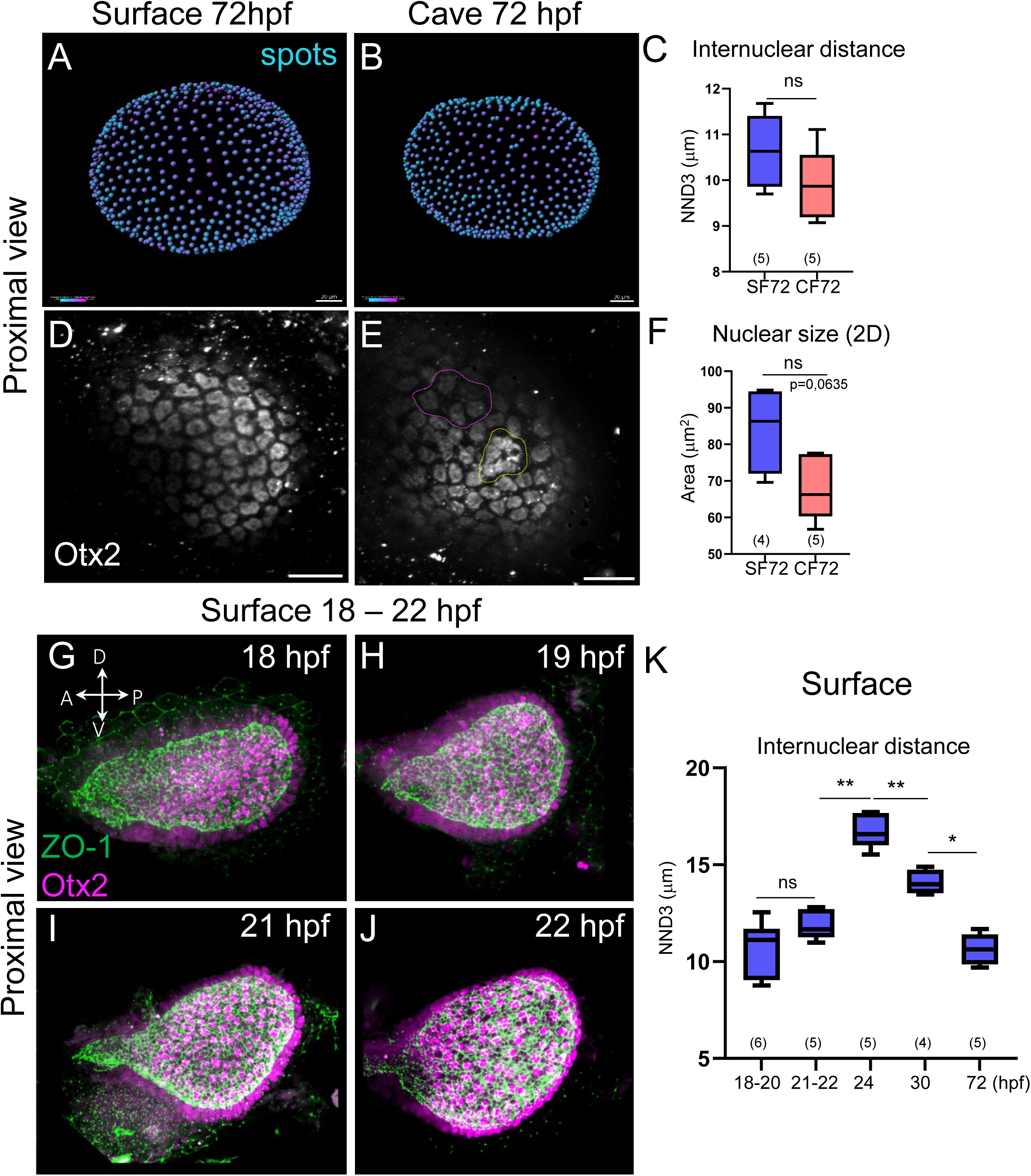
Temporal dynamics of RPE stretching reveal transient expansion of proximal cells during optic cup morphogenesis. (A,B) Colour-coded 3D maps of NND3 across the RPE surface for SF and CF at 72 hpf. At this stage, proximal RPE nuclei, shown here as dots, display compact organization in both morphs, indicating completion of RPE spreading. (C) Quantification of mean NND3 in the proximal RPE. NND3 values are not statistically different between the two morphs. (D,E) Proximal views of the OC from both morphs at the same stage, showing the organization of nuclei immunostained for Otx2 (grey). The magenta and yellow dotted line in (E) delineate irregular clustering compare to more regular nuclear spacing in SF (D). (F) Quantification of nuclear surface area in the proximal RPE. (G–J) Proximal views of dissected OC from SF at the indicated stages, immunostained for Otx2 (magenta) and ZO-1 (green). The temporal sequence illustrates the change in nuclei organization and OC shape. (K) Temporal analysis of proximal RPE NND3 from 18 to 72 hpf. A sharp increase is observed between 21–24 hpf followed by a progressive decrease after a peak at 24 hpf. Scale bars: 100 µm (A–B), 20 µm (D–E).

To characterize the temporal dynamics of RPE stretching in SF, we also examined both the early (between 18 and 22 hpf) and late (30hpf) stages of OC morphogenesis. We observed that the onset of RPE stretching occurs in parallel with the elongation and the change in shape of the optic vesicle (Figure 4G-J and Fig. S6). Quantification of NND3 revealed a rapid increase between 21 and 24 hpf (Figure 4K). This corresponded to a peak in NND3 values, which then decreased. These results suggest that the proximal RPE stretches only during the period of OC folding.

Finally, intrigued by the lower NND3 values observed in the ventral region of the distal RPE at 24 hpf compared to the dorsal region (Figure 3A’,H), we analysed ventral nuclear organization at 36 hpf in CF. At this stage, OC morphogenesis was still incomplete (Figure 1J) and Otx2 expression was downregulated in the RNE. This made it easier to analyse RPE anatomy in this region. Quantification of NND3 in CF at 36 hpf in the distal region showed that the spacing of the ventral nuclei was greater than at 24 and 30 hpf in the dorsal region (Figure 3E, Movie 7, and data not shown). Taking NND3 as a proxy of stretching, these data demonstrate that the timing of stretching in the distal RPE is regionally controlled, with ventral regions stretching much later than dorsal regions. This occurs at a time when the proximal cells have already ceased their stretching and begun to re-constrict (Figure 4K).

### The most distal cells of the RPE are small, elongated and aligned along the OC meridians

The NND3 measurements above indirectly suggested significant cell size heterogeneity within the RPE. To compare cell sizes between the two regions quantitatively, we implemented complementary strategies (Figure 5AB,EF). In the distal RPE, accurate segmentation of individual cell outlines was not possible due to the close apposition of apical RPE and RNE membranes and the discontinuous ZO-1 signal (Figure 5B’; see Materials and Methods). We therefore used nuclei-derived centroids to compute Voronoi surfaces (Aurenhammer, 1991) and used it as a proxy for cell surface area (Figure 5C). While Voronoi tessellations do not reflect true cell contours (Kaliman et al., 2016), they provide a robust estimate of relative cell surface area across a large population of cells. For proximal RPE cells, we manually segmented the cell outlines at a slightly earlier stage than 24 hpf (Figure 5G and Fig S7) as the close proximity of the two layers and weak ZO-1 signals in the RPE cells (Figure 2D) made it impossible to resolve the cell borders. Using this approach, we found that the apical surface area of distal RPE cells was approximately one-third that of proximal cells, which exhibited irregular polygonal shapes with a mean area of 280 ± 70 µm² (Figure 5D, FG). Similarly, measurements of nuclear surface areas in 2D projections in SF at 24 hpf revealed smaller nuclei in distal than in proximal RPE regions (Figure 5H). These data confirmed the heterogeneity of cell size within the RPE, which further strengthens the emergence of a stretching gradient along the proximo-distal axis.

**Figure 5.**
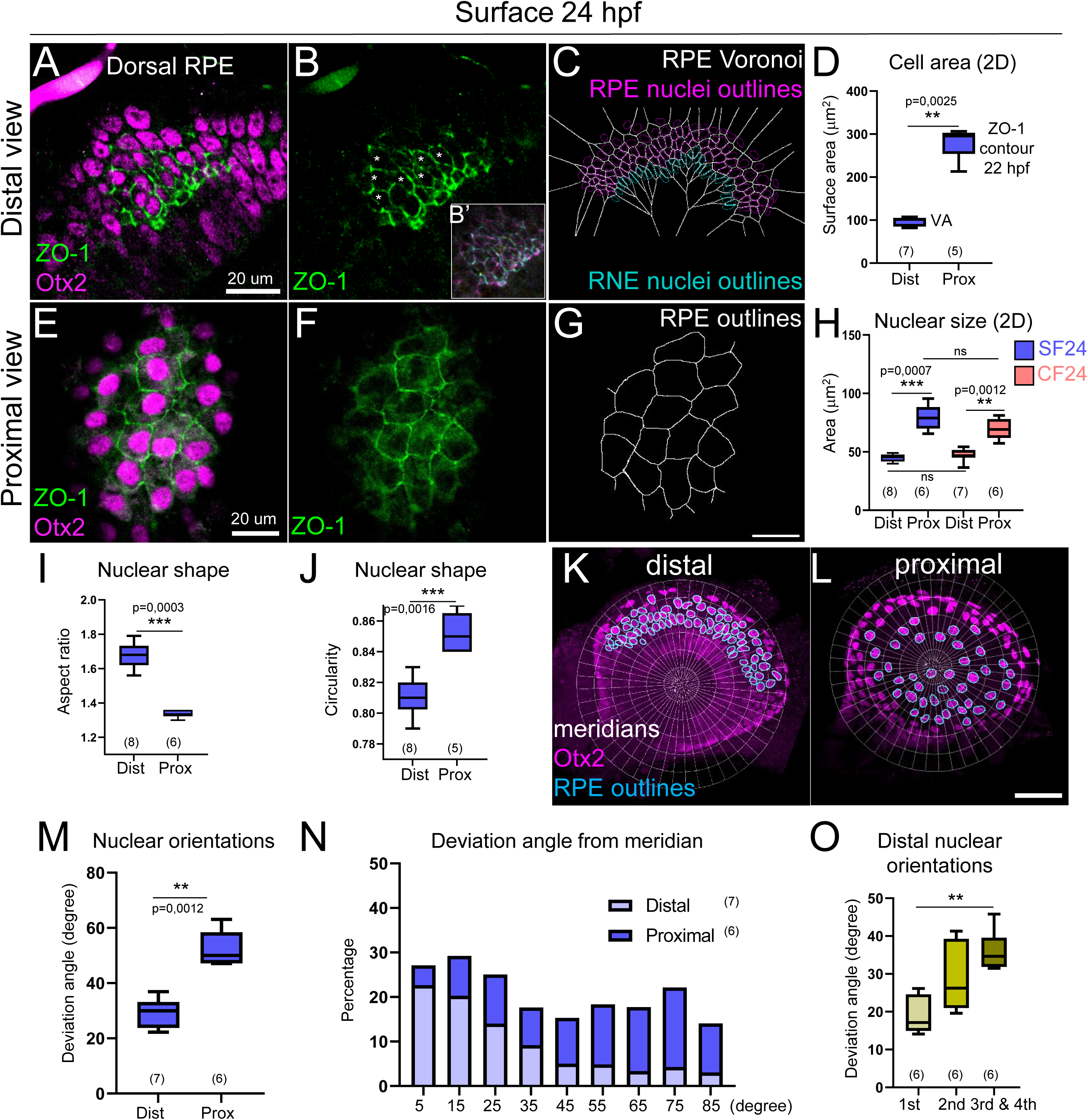
Distal RPE cells display elongated morphology and preferential alignment along optic cup meridians. (A,B,E,F) Maximum intensity projections of distal and proximal RPE at 24 hpf in SF. Dissected OC were immunostained for ZO-1 (green) and Otx2 (magenta). ZO-1 accumulation outlines small and elongated distal RPE cells (B, asterisks) with straight apical borders and bigger cells in the proximal region, with no such features (F). The inset in panel B (B’) shows depth-colour-coded maximum-intensity projection that highlights the close proximity of the apical sides of the two epithelia. (C) Voronoi surface (grey) generated from nuclei-derived centroids for quantification of apical surface areas. The RPE nuclei outlines are in magenta. The uppermost distal row of RNE nuclei appears in turquoise. (G) RPE cell contours of proximal RPE cells. (D) Quantification of apical surface areas for distal and proximal RPE cells (calculated from C and G). (H) Quantification of nuclear size (surface area) for the two morphs at the indicated stages. (I,J) Quantification of nuclear shape descriptors showing aspect ratio (I) and circularity (J) in distal and proximal nuclei. (K,L) Distal and proximal maximum intensity projections merged with the reference meridian map used to analyse nuclear orientation from RPE nuclei outlines. (M-O) Quantification of nuclear alignment to the meridians of the reference map in the distal and proximal regions of the RPE, expressed as a deviation angles. Scale bar: 20 µm (A-C, E-G), 50 µm (K,L).

The markedly reduced stretching and the compact organization of distal RPE nuclei near the ventricle margin suggested that these cells might contribute differently to the tissue expansion. This prompted us to examine the shapes and orientations of RPE cells. At 24 hpf in SF, ZO-1 accumulation in the distal RPE revealed elongated cells with straight borders, suggesting mechanical tension in cell junctions (Figure 5AB). In contrast, no such features was observed in proximal cells (Figure 5EF).

In order to quantify the differences in nuclear shape between the proximal and distal regions of the RPE, we measured two parameters: aspect ratio and circularity. As shown in Figure 5IJ, distal nuclei exhibited higher aspect ratios, consistent with elongation, whereas proximal nuclei displayed higher circularity values, indicative of a more rounded 2D shape (Movie 8) and suggesting a lack of preferential orientation. To assess nuclear orientation, we overlaid a reference map of OC meridians aligned to the lens center and analysed the first four rows of nuclei (Figure 5KL and see Materials and Methods). For each nucleus, the angle of its longest axis was compared to the closest meridian, and the deviation between both angles was computed as a measure of alignment. Distal RPE nuclei showed a two-fold lower mean deviation angle than proximal nuclei (Figure 5M). Notably, over 40% of distal nuclei exhibited a deviation angle below 20 (Figure 5N), whereas proximal cells displayed no preferred orientation (Figure 5L-N). Furthermore, we found that the nuclei in the most distal rows displayed a substantially lower deviation angle than the nuclei in the third and fourth rows (Figure 5O).

Together, these analyses demonstrate that distal RPE cells differ markedly from proximal cells in size, shape and orientation. They exhibit preferential, graded alignment and elongation along OC meridians and seem to have straighter apical borders. This cellular organization strongly suggests that the most distal RPE cells are under tension, allowing cells to resist deformation while accommodating mechanical constraints at the OC margin.

### Nuclear volume gradient within the RPE

Numerous studies have observed consistent relationships between a cell’s DNA content, nuclear size, and cell size across unicellular and multicellular eukaryotic species (Balachandra et al., 2022). Having found that the apical surfaces and nuclear areas of distal cells of the RPE were smaller than those of proximal cells (Figures 5D and 5H), we wondered whether their volumes differed, too. To address this, we used the Otx2 signal to segment RPE nuclei in 3D with the IMARIS software and analysed the volumes from generated surfaces (Figure 6AB). Strikingly, the volume of proximal nuclei were 2.5 fold larger than distal nuclei at 24hpf in SF (Figure 6C), which was consistent with the 2D measurements shown above for the two morphs (Figure 5H). Because automatic segmentation based on Otx2 signal intensity may introduce bias into nuclear size estimation, we performed an independent validation using manual segmentation. Using the DAPI signal, we manually outlined 6-8 RPE nuclei per sample across the proximal regions (Figure 6E’ and Fig. S9) and confirmed that proximal RPE nuclei are larger in volume than distal ones in both morphs (Figure 6CD and data not shown). These results strongly suggest that changes in cell shape associated with stretching are proportional to an increase in 3D cell size.

**Figure 6:**
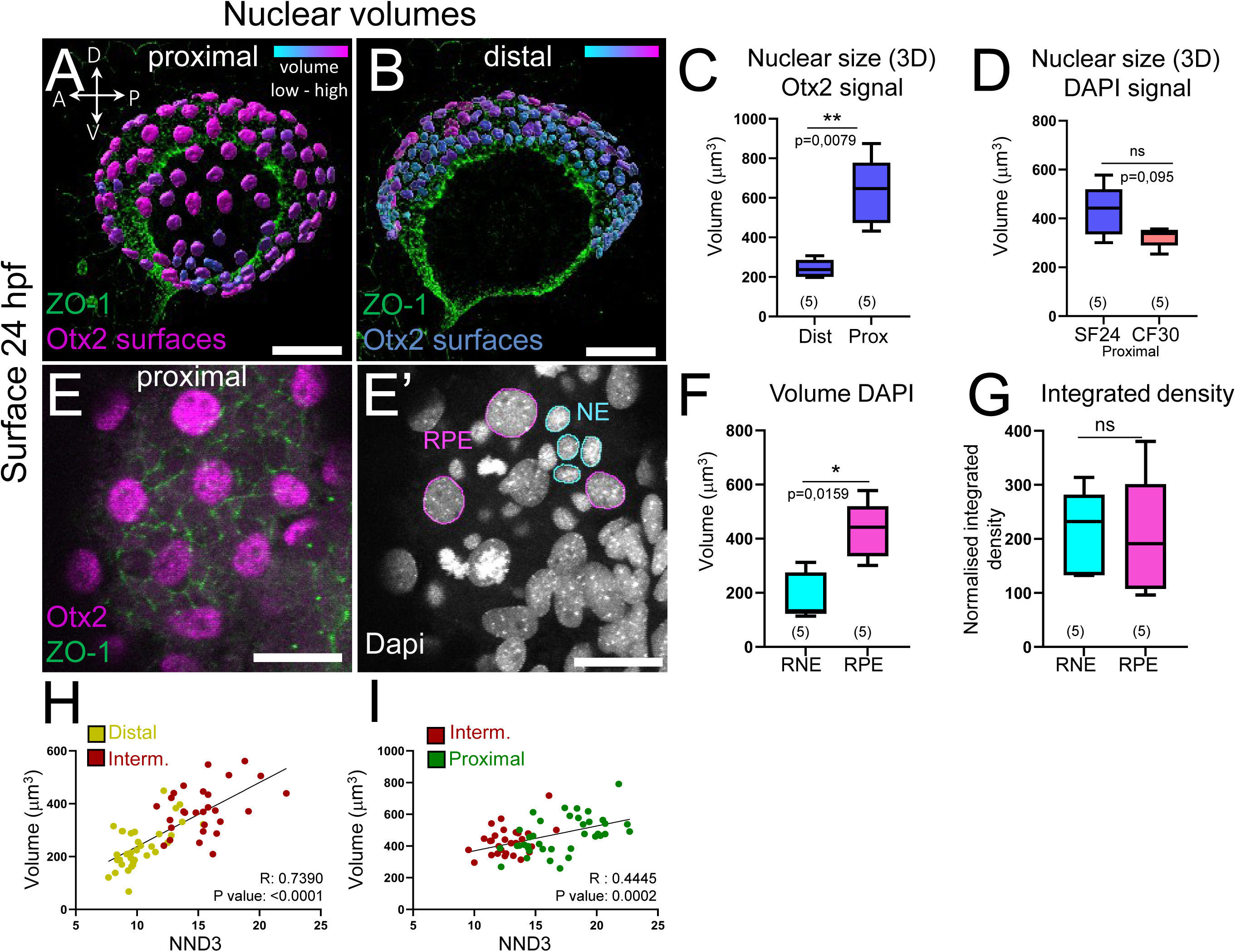
Cell size gradient along the proximo distal axis. (A,B) 3D renditions of Otx2-labelled RPE nuclear surfaces from SF embryos at 24 hpf, showing the proximal and distal regions respectively. The colour code, which ranges from low to high value, highlights the differential 3D size of RPE nuclei. (C) Quantification of mean nuclear volumes in distal and proximal RPE regions of SF. (D) Quantification of nuclear volume in the proximal region of the RPE in SF and CF at equivalent stages of OC morphogenesis. (E,E’) Maximum intensity projections of the proximal RPE from a SF OC immunostained for Otx2 and ZO1 (magenta and green in E) and counterstained with DAPI (grey in E’) for DNA content quantification. (F-G) Quantification of nuclear volumes and integrated DAPI fluorescence intensity in proximal RPE and retinal progenitors. Scale bars: 100 µm (A,B), 50 µm (E,E’). (H-I) Distribution of nuclei volume according to their respective NND3. The R coefficient (linear regression) are indicated (G) Temporal increase in the volume of distal RPE nuclei during OC folding in SF. The width, height and length of each nucleus were measured manually for five RPE nuclei at three different times. These measurements were then used to calculate the volumes of the nuclei using the ellipsoid formula.

To determine whether nuclear size followed a gradient along the proximo-distal axis, we also compared volumes (using Otx2 signal) in the distal, intermediate, and proximal regions (as performed for the NND3 parameter in Figure 3L). Nuclear size increased progressively from distal-to-proximal regions (Figure 6AB and Fig. S8). Moreover, analysis of successive distal RPE ‘rows’ revealed a pattern within the distal domain itself: the most distal nuclei were approximately half the size of those in the third and fourth rows (Fig. S8).

In addition, we found a significant positive correlation (R = 0.7390 and 0.4559, Pearson coefficient) between the volume of nuclei and their respective NND3 in different regions of the RPE (Fig. 6HI). This result shows a relation of proportionality between RPE nuclear size and stretching, regardless of the region or extent of cellular stretching.

Together, these results further demonstrate that during the second phase of RPE expansion, RPE cells undergo differential changes in both shape and growth. Noteworthy, they exhibit graded nuclear enlargement along the proximo-distal axis, proportional to the extent of regional stretching.

A process commonly used for increasing cell volume is to increase DNA content through endoreplication (Balachandra et al., 2022), (Øvrebø and Edgar, 2018). To test whether the increase in nuclei volume in proximal RPE cells occurs through endoreplication, we segmented nuclei manually in 3D using the DAPI signal in two cell types (RPE cells and retinal progenitors) in the proximal region of the OC (Figure 6EE’ and Fig. S9). RPE nuclei were 2.5-fold larger than retinal progenitor nuclei (Figure 6F) and displayed reduced mean DAPI intensity (data not shown). However, the integrated DAPI fluorescence intensity did not differ between the two cell populations (Figure 6G). This showed that nuclear enlargement in proximal RPE cells is not associated with increased DNA content and does not result from endoreplication, but may rather reflect changes in nuclear organization, chromatin architecture, or transcriptional activity.

### Proximal RPE stretching and OC shape and size

Since the amplitude of proximal RPE deformation at 24 hpf was lower in CF than in SF, and was more variable in CF at 30 hpf than in SF at 24 hpf (Figures 3A–D), we hypothesized that proximal stretching follows distinct scaling dynamics in CF. We first quantified cell surface expansion in this region. We estimated cell surface areas using Voronoi tessellation (Figures 7A-D). As expected, this parameter was lower in CF than in SF at 24 hpf (Figure 7E), confirming a reduced degree of stretching at this stage, consistent with the lower NND3 values (Figure 3D). In CF, the Voronoi surface area at 30 hpf was increased compared to 24hpf (Figure 7E), and was similar to SF at 24 hpf, also in agreement with NND3 values (Figure 3D). By contrast, the geometry of the Voronoi surfaces in the proximal region differed between the two morphs (Figure 7F), consistent with the greater variability in nuclear positioning in CF (Figure 7CD) and the higher variance in NND3 values (Figure 3D). While the proximal nuclei were more circular in SF at 24hpf (Figure 5JL and Figure 7A), they exhibited abnormal shapes in CF at 30 hpf, an “equivalent” stage of RPE morphogenesis (Figure 7C and data not shown).

**Figure 7:**
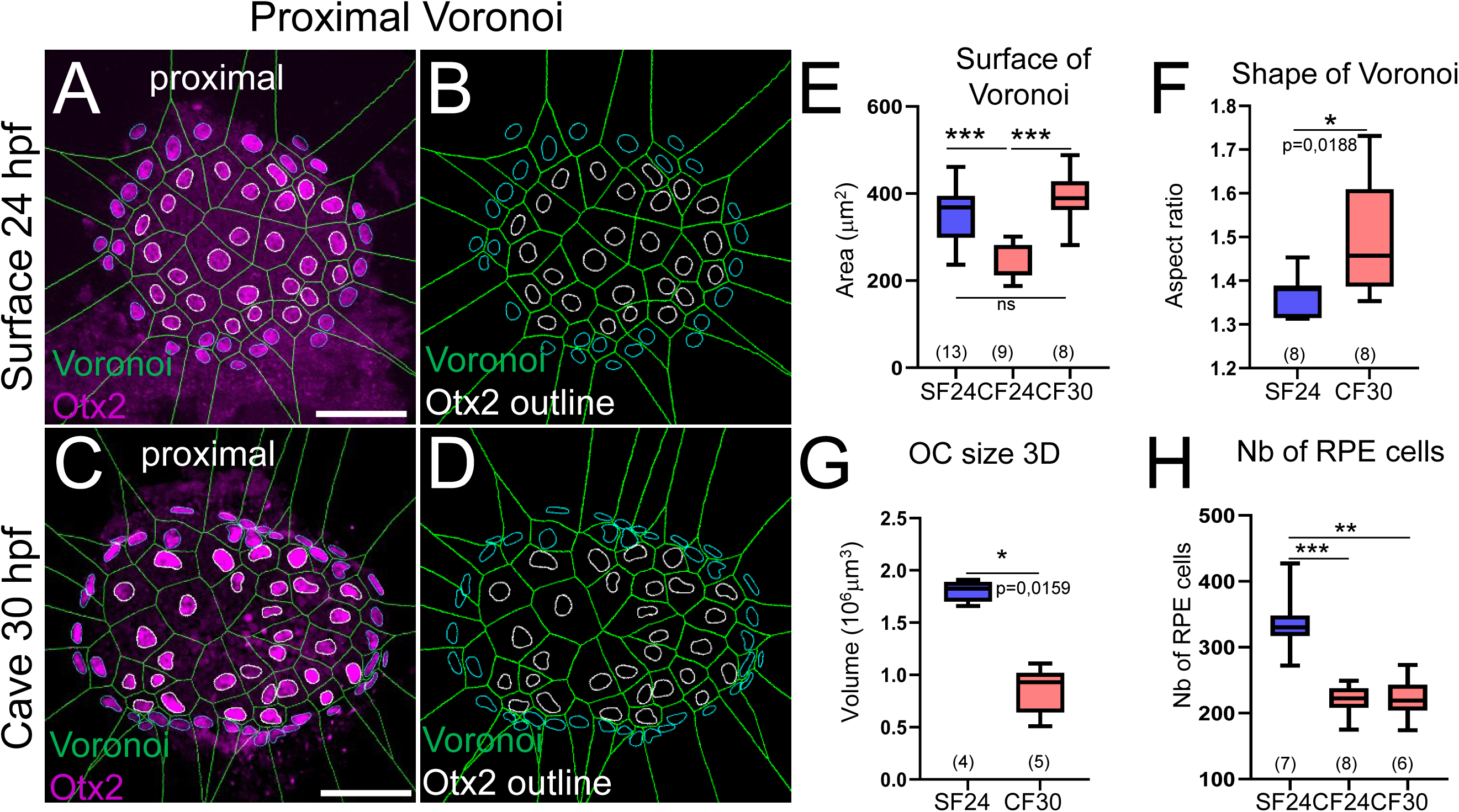
Proximal RPE stretching, OC size and number of RPE cells. (A-D) Merges combining RPE nuclear outlines and derived proximal Voronoi maps, as well as Otx2 staining (A,C), were used to analyse the proximal region in SF and CF at an equivalent stage of OC morphogenesis. White nuclear outlines were used for the quantifications. (E) Quantification of mean Voronoi surface areas in proximal RPE in the two morphs at the indicated stages. (F) Quantification of Voronoi surface aspect ratios in the proximal RPE in SF and CF at the indicated stages. (G) Quantification of the volume size of the OC of the two morphs at the indicated stages. (H) Quantification of the total number of RPE nuclei in the two morphs at the indicated stages. Scale bars: 20 µm.

In a second approach, we compared the size of the OC and the total number of RPE cells in the two morphs (Figure 7GH). The OC was twofold smaller in CF at 30 hpf than in SF at 24 hpf (Figure 7G); the total number of RPE cells was 1.5-fold lower in CF at both stages (Figure 7H); and the Voronoi surface area of proximal RPE cells in CF at 30 hpf was comparable to that of SF at 24hpf (Figure 7F).

Together these data show that RPE number of cells, proximal stretching and RNE size are scaled in the CF OC, which may be initially due to the proportional allocation of RPE and RNE territories at the stage of RPE cell specification.

## Discussion

### The expansion of the RPE occurs in a spatio-temporal pattern

Our study provides a detailed characterisation of the RPE during its expansion phase during OC morphogenesis in a teleost fish. By combining quantitative analyses of ventricular edge progression and spatial organization of RPE nuclei and cell morphology, we demonstrate that RPE expansion is not uniform (Figure 8) and found that dorsal and ventral distal RPE expand asynchronously. On the contrary, RPE expansion follows a spatio-temporal pattern of anisotropic deformation, exhibiting a differential mechanical regime along the proximo-distal axis. This process is transient, suggesting that RPE stretching is restricted to a specific morphogenetic window and likely reflects the temporal dynamics of mechanical forces acting during OC folding. The fact that proximal RPE stretching ceases after OC folding suggests that this deformation is closely coupled to evolving tissue constraints, rather than being a continuous intrinsic process.

**Figure 8.**
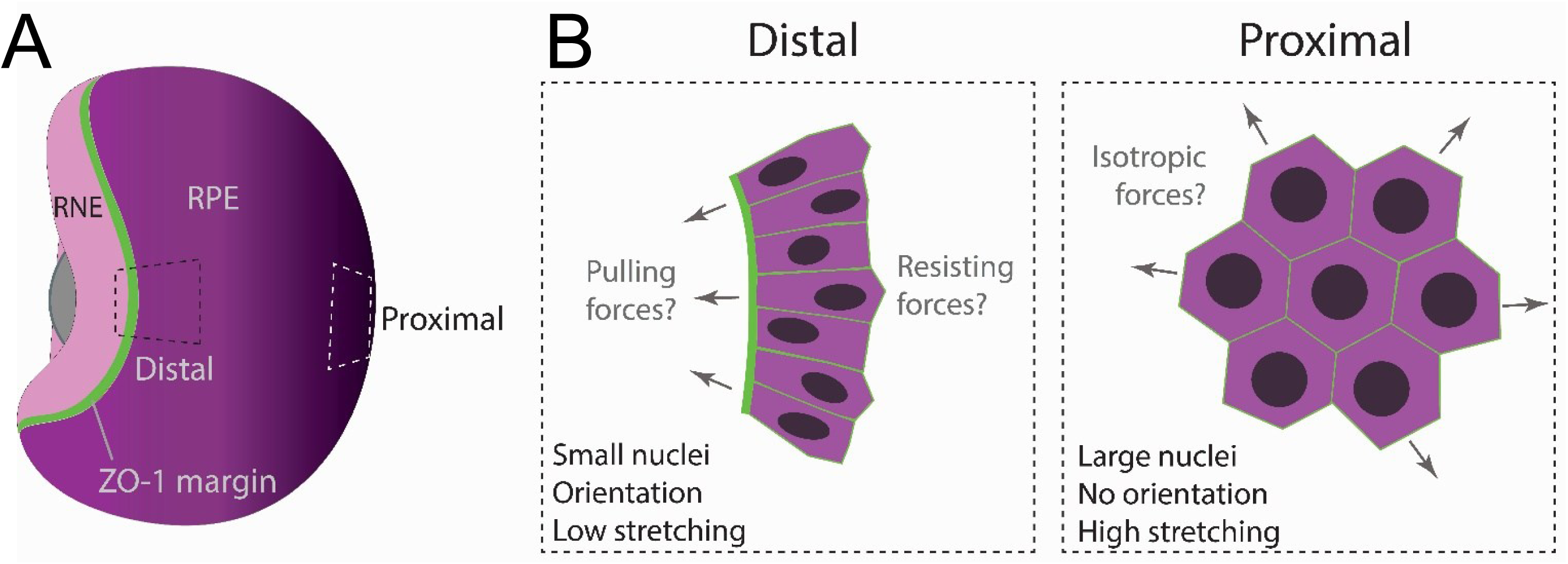
Emergence of nuclear size and cellular stretching gradients in the RPE during OC folding. **(A)** 3D drawing in a slightly tilted dorsal view of a 24hpf OC. The external view represents the RPE as it spreads during OC folding. The most distal circumferential edge of the ventricle (the ZO-1 margin), marks the boundary between RPE and the RNE cells. The colour gradient for the RPE illustrates the gradients in cellular morphology along the proximal-distal axis. (B) The diagrams depict the distinct morphologies of RPE cells in the distal and proximal regions. These cells differ from each other in terms of size, shape, orientation and stretching. Hypotheses regarding putative in plane forces at play are indicated. Orthogonal out-of-plane putative forces emanating from the bulging RNE and that would participate in compression force onto the RPE are not indicated.

Previous work in zebrafish described tissue movement directionalities during RPE spreading, based on marker expression and live imaging analyses (Kwan et al., 2012, Cechmanek and McFarlane, 2017, Moreno-Mármol et al., 2021). Our results in *Astyanax mexicanus* reveal a graded and sequential stretching behaviour, in which proximal RPE cells undergo early and pronounced isotropic extension, while distal cells remain compact initially during OC folding and stretch later in a coordinated fashion.

### Cell stretching as a conserved morphogenetic strategy

Stretching-mediated epithelial morphogenesis represents a conserved morphogenetic strategy across metazoans, associated with the emergence of cell morphological asymmetries within initially homogeneous epithelia. In the mammalian blastocyst, lumen expansion generates hydrostatic pressure that drives stretching and flattening of the trophectoderm (Chan and Hiiragi, 2020). In the *Drosophila* wing imaginal disc, peripodial membrane epithelial cells undergo a cuboidal-to-squamous transition in response to tissue-extrinsic forces, leading to emergence of gradients in cellular morphology (Harmansa and Lecuit, 2024). In the adult ovary, follicle cell stretching is tightly coordinated with transcriptional changes during morphogenesis (Jia et al., 2022). Similarly, studies in zebrafish showed that RPE cells undergo epithelial flattening during OC formation (Cechmanek and McFarlane, 2017; Moreno-Mármol et al., 2021), a process coupled to transcriptional reprogramming (Buono et al., 2021).

Our findings on the species *Astyanax mexicanus* build on and add to this framework, integrating an evo-devo approach that compares different morphs within the same species. The gradients in nuclear and cellular morphology along the proximal-distal and dorso-ventral axes of the RPE appear to be proportional to the degree of stretching (Figure 8). We show that this RPE expansion morphogenetic program exhibits evolutionary plasticity, with altered spatial and temporal coordination in CF. Thus, while conserved as a morphogenetic strategy, the spatial and temporal patterning of RPE cell stretching would “follow” developmental evolution of the OC on a micro evolutionary scale. Because OC morphogenesis is conserved among teleosts (Cardozo et al., 2023), conducting comparable quantitative analyses in zebrafish and medaka will enable to test whether spatially and temporally patterned asymmetries of the RPE constitute a conserved morphogenetic module at a macro evolutionary scale.

### Mechanical drivers of proximo-distal asymmetry in RPE stretching

The graded stretching along the proximo-distal RPE axis suggests differential distribution of external forces and/or spatially regulated mechanical properties within the RPE, or both. Direct measurement of forces *in vivo* remains challenging in *Astyanax*, but several observations suggest a potential combination of in-plane forces acting differentially along the tissue (Figure 8), which could be more easily tested in zebrafish and its associated genetic tools.

By analogy with lumen-driven deformation in the mammalian blastocyst (Chan and Hiiragi, 2020), the bending of the RNE as seen in live imaging movies of zebrafish (Heermann et al., 2015; Nicolás-Pérez et al., 2016) and *Astyanax* (data not shown and Devos et al. 2021), could locally increase intraventricular pressure, exerting perpendicular mechanical compression on the adjacent RPE, primarily on the proximal region. In line with this hypothesis, we observed an increased intraventricular space in CF at 24 hpf compared to SF at the same stage (Fig. S1 and data not shown).

We also observed major inter-individual variability in OC size between CF samples at 30 hpf, with volumes ranging from 0.5 to 1.1.10^6^ µm³ (Figure 7G). Interestingly, a twofold variation in the size of intraventricular space was also detected between CF samples (data not shown), suggesting a possible correlation between RNE size and the extent of intraventricular space reduction. Notably, further using the high phenotypic variability within CF or analysing second generation SFxCF hybrids could enable testing whether OC size, the number of RPE cells, the intraventricular width, and the NND3 co-vary. This would provide further support for a link between RNE growth and RPE stretching.

At the same time, the circumferential epithelial flow of the RNE (rim movement) generates tangential forces at the OC margin (Kwan et al., 2012; Heermann et al., 2015). Defects in OC folding are more severe when rim cell movement is impaired than when basal constriction of the RNE alone is disrupted (Sidhaye & Norden 2017). We hypothesize that this flow could exert tensions on most distal RPE cells, resulting in their elongated shape and preferential alignment along meridians. The most distal RPE would then function as a mechanically reinforced domain that “resists” early deformation, maintaining tissue integrity during folding before undergoing delayed stretching. However, we cannot rule out that the most distal RPE cells could also behave like “leader cells” during collective migration. It would be interesting to investigate whether they generate basal protrusions (such as lamellipodia or filopodia) to propel themselves forward and gradually draw the RPE behind them.

Together, these observations strongly suggest a model in which proximal and distal RPE cells experience distinct mechanical regimes, arising from their relative position within the deforming OC and their coupling to the RNE tissue dynamics.

### Active versus adaptive roles of the RPE in OC folding

Several independent studies have shown that, in fast developing species, RPE expansion during OC morphogenesis relies primarily on cell shape changes rather than cell proliferation (Kwan et al., 2012; Moreno-Mármol et al., 2021). However, whether the RPE functions as a passive epithelium adapting to forces generated by the RNE, or instead plays an active mechanical role in shaping the OC, remains debated. It was initially speculated that the RPE massively enlarging its surface could generate a force for the epithelial flow (Heermann et al., 2015). Targeted disruption of cytoskeletal dynamics within the RPE impairs OC folding, supporting the idea that RPE stretching actively contributes to OC morphogenesis (Moreno-Mármol et al., 2021). Consistently, a recent physical modelling study suggests that active in-plane collective behaviours at the apical surfaces of both the RPE and the RNE generate patterned spontaneous strain sufficient to drive tissue curvature and RNE basal invagination (Ramos et al. 2025). The compact organization of small, elongated distal RPE cells aligned along meridians and accumulating high levels of tight-junction markers, as observed in our study, suggests that the RPE may act as a mechanical interface transmitting resisting forces from its most distal rows. Based on these observations, we propose a working model in which the proximal RPE stretching results from a balance between two opposing mechanical inputs: (1) compression forces exerted by the massive and highly mitotic retinal RNE, and (2) distal resisting forces generated by the tightly connected RPE cells. In this framework, it is tempting to speculate that the deformation of proximal RPE cells could arise from the interplay between such compressive and tensile cues. Laser ablation experiment in zebrafish to disrupt distal RPE junctions or RNE integrity will help assess the potential contributions of these forces to proximal RPE stretching.

### Nuclear size gradients correlate with nuclear spacing gradients

A striking feature of RPE morphogenesis revealed by our study is the pronounced gradient in nuclear size along the proximo-distal axis (Figure 8). Proximal RPE nuclei are two to threefold larger in volume than distal ones, and this variation correlates with gradients in cell stretching and internuclear spacing. Consistent with this spatial gradient, direct measurements indicate that RPE nuclei also grow over time during OC folding, supporting the idea that nuclear enlargement is dynamically regulated during morphogenesis. Given that nuclear volume and cytoplasmic volume are tightly correlated (Balachandra et al., 2022), this strongly suggests that the actual volume of RPE cells follows the same gradient as nuclei. Polyploidization offers cells several potential fitness benefits, including the ability to increase cell size and biomass production (Øvrebø and Edgar, 2018). However, our measurements of DNA content argue against endoreplication as the primary cause of nuclear enlargement. In this context, the marked nuclear size increase in proximal RPE cells could represent an adaptive response that could support increased metabolic or transcriptional demands associated with transient cell deformation. Interestingly, several cellular processes involved in nuclear size control, including nucleocytoplasmic transport, LINC complexes, RNA processing, regulation of nuclear envelope expansion and partitioning of importin α, have been implicated in coordinating nuclear growth with cellular state (Cantwell and Nurse, 2019). Mechanical forces are known to influence nuclear architecture, transcriptional activity, and cell-cycle dynamics (Lyer et al., 2012; Vivo et al., 2024). It is therefore tempting to hypothesize that the proximo-distal gradient in nuclear size may also reflect active, mechanotransduction-related modulation of nuclear function and cell state. Future analyses combining cell-cycle markers, transcriptional readouts, and mechanotransduction reporters will be required to test these hypotheses.

### Regionalization and patterning of the RPE

Recent transcriptomic studies in primates demonstrated that the mature RPE exhibits continuous, location-dependent gene expression variations despite relatively subtle morphological differences (Mungale et al., 2022). These data challenge the view of the RPE as a homogeneous epithelium and instead support the existence of spatially distinct cellular states and functions. Our study in *Astyanax* suggests mechanical regionalization of the RPE during its morphogenesis. Gradients in cell stretching, nuclear size, and epithelial organization indicate that distinct RPE domains experience different physical constraints and may contribute differentially to OC morphogenesis. Such mechanical heterogeneity could affect early molecular regionalization within the tissue. Interestingly, pigment-associated genes are first expressed in a restricted dorso domain and maintain spatial heterogeneity throughout OC formation (Cechmanek and McFarlane, 2017), raising the possibility that early molecular patterning could also prefigure later mechanical behaviours. Conserved axial patterning pathways that establish dorsal-ventral and nasal-temporal identities in the retina, such as BMP signalling, also act in non-neural ocular tissues (French et al., 2009; Erickson et al., 2010). Future studies linking molecular regionalization to mechanical heterogeneity in the RPE will address whether cell stretching is related to spatial position or cell identity.

### Cavefish as a window into morphogenetic robustness and evolution

The cave-adapted morph of *Astyanax mexicanus* provides a natural experiment for studying the robustness and evolutionary plasticity of OC morphogenesis. The OC in CF are smaller and exhibit delayed folding compared to SF embryos, yet they ultimately achieve complete invagination together with RPE expansion (Devos et al., 2021). This heterochrony reveals the temporal flexibility of the morphogenetic program. Furthermore, the reduction in size of both the RNE and the RPE in CF, as well as the stretching of proximal cells of a similar magnitude to that observed in SF, further suggest that mechanical constraints and/or intrinsic growth regulation play a critical role in coordinating the morphogenesis of the two adjacent tissue layers. Inter-individual variability in CF will be useful for (de)correlating the size and shape of the OC with abnormalities in RPE cell stretching. Finally, the differences shown here between the two eco-morphs of *Astyanax* highlight the extent to which significant changes in morphogenesis can occur over short evolutionary timescales (Fumey et al., 2018; Herman et al., 2018; Policarpo et al., 2024). In the CF, the RPE exhibits early defects in nuclear shape and positioning at 30 hpf, which result into abnormal RPE nuclei clustering at 72 hpf. These alterations to RPE morphogenesis in CF could then have important secondary functional consequences as indicated by the reduced ONL width observed at 72 hpf (Fig. S5), and the absence of photoreceptor outer segments (Emam et al., 2020). In the dual model of ocular degeneration in CF, both the lens and the RPE may contribute to the onset and induction of progressive vision loss in CF (Ma et al., 2020). Such defective function of the CF RPE could stem, at least in part, from impaired morphogenesis, patterning and final differentiation.

## Conclusions

Overall, our study reveals that RPE morphogenesis during OC folding in fish is governed by a temporally and regionally patterned sequence of cellular behaviours. Proximal RPE cells stretch the most, while the most distal cells maintain compactness and alignment along meridians. This pattern suggests a dual model in which the distal RPE would ensure a resisting force through tension balance along the circumferential margin while the proximal cells would experience a tangential, out-of-plane-force (from the bulging RNE) that would contribute to their stretching. What are the intrinsic mechanisms allowing RPE cells to stretch? Moreno-Marmol et al. (2021) showed that localized interference with the RPE cytoskeleton disrupts tissue stretching autonomously in zebrafish. Here, in *Astyanax* we observed an increase in nuclear size without endoreplication accompanying the stretching of proximal cells, which suggests a causal link.

## Material and methods

### A. mexicanus strains

Our colony of surface and cavefish originated from parents obtained from William Jeffery’s laboratory in 2004. The surface fish come from rivers in Texas and the cavefish come from the Pachón cave in the Mexican state of Tamaulipas. We obtained the embryos of the two morphs through *in vitro* fertilization after inducing maturation and reproduction in the breeding colony by altering the water temperature. The embryonic development of *A. mexicanus* at 24°C is similar and synchronous for both morphs (Hinaux et al., 2011). We raised and treated animals in accordance with the French and European regulations that govern the use of animals in research. Authorization for the use of animals in research is 91-116. As all experiments were performed on embryos at 18-72 hpf, an authorization from the “Comité d’éthique en matière d’expérimentation animale Paris Centre et Sud” was not required.

#### Embryo staging and fixation

The embryos were staged using morphological criteria and their known developmental times (Hinaux et al., 2011). The embryos were fixed in 4% paraformaldehyde in PBS, dehydrated using a series of ethanol/PBS solutions and stored in ethanol at −20 °C.

#### Immunofluorescence and antibodies

The fixed embryos were rehydrated through a series of graded ethanol/phosphate-buffered saline (PBST) solutions before being washed in PBST. The samples were depigmented for 5-10 minutes for 24-30 hpf embryos or up to 20 minutes for 40-72 hpf embryos in a 5% H₂O₂ solution in PBST under intense light. After washing, antigen retrieval was performed by incubating the samples in Histo-VT at 68 °C for one hour, followed by several washes in PBST. The embryos were then bathed in a blocking solution containing 10% goat serum, 1% Triton X-100 and 1% dimethyl sulfoxide in PBST at room temperature for two hours. The samples were then incubated with the following primary antibodies: rabbit anti-Otx2 (Abcam; ab183951, cat# EPR20375) and mouse anti-ZO-1 (Thermo Fisher Scientific, clone 1A12, cat# 33-91100) at a dilution of 1:500 in blocking solution at 4°C for up to two days. After washing in PBST, the samples were incubated with Alexa Fluor-conjugated secondary antibodies (1:1000) and DAPI (Sigma, cat# 10236276001) for two days at 4 °C in the dark. Finally, the samples were washed in PBST at 4 °C.

#### Dissection of the OCs and mounting

The procedure was performed in two steps under a binocular fluorescence microscope in a dark room at room temperature. Ultraviolet lighting was used to visualise the DAPI labelling. The OC dissection was performed in a Petri dish covered with a thin layer of transparent agar and containing PBST to facilitate handling of the embryos. The yolk sacs and the region posterior to the otic vesicle were removed using tungsten needles. The heads were transferred to a glass slide using forceps. The PBST was removed before being replaced with 10-15 µl of anti-fading mounting medium (VECTASHIELD). The OCs were separated from the brain using tungsten needles and any residual brain fragments were carefully removed, except for the olfactory epithelium. Two OCs were placed in the centre of an eyelet containing 10 µl of mounting medium, oriented either in proximal or distal view. The preparation was then covered with a coverslip and sealed using nail varnish.

#### Fluorescent image stack acquisition

Confocal image stacks were acquired using a Leica SP8 confocal microscope equipped with Leica Application Suite software. A water-immersion HC PL APO 40×/1.10 W CORR CS2 objective was used for most experiments, while a HC FLUOTAR L 25×/0.95 W VISIR objective was employed for samples older than 30 hpf. All images were collected at a bidirectional scan speed of 400 Hz. Laser power and photomultiplier gain were optimized individually for each sample and channel, and no Z-axis compensation was applied. Channels were acquired sequentially with emission ranges adjusted to prevent signal bleed-through. For samples imaged at 40×, image stacks typically comprised 70-100 z-steps for half OCs (oriented either distally or medially) and about 200 z-steps for entire OCs. At 40× magnification, voxel size was 0.24 µm × 0.24 µm × 0.42 µm (x-y-z), and 0.6 µm × 0.6 µm × 1 µm (x-y-z) at 25x. Images were recorded in 8-bit format at a resolution of 1024 × 1024 pixels (40×) or 512 × 512 pixels (25×).

#### Image processing

Raw data image processing was performed in Fiji/ImageJ software. Quantifications were performed in Fiji/ImageJ or Imaris 10 (Bitplane) softwares.

#### Distance from ZO-1 margin to lens and Distal ZO-1 surface area

Maximum intensity projections of the distal region were used to measure (i) the distance (d) between the distal margin of the ZO-1 labelling and the outer edge of the lens, and (ii) the distal surface area (S), defined as the projected area covered by ZO-1 labelling. Although this 2D projection-based approach does not capture exact 3D geometry, it provided a robust and reproducible proxy for RPE expansion. The distance d was measured manually using the Line tool in Fiji, and S was quantified manually using the Polygon tool and ROI Manager.

#### Nearest neighbour distances in 3D (NND3)

Related to Figure 3, Figure S3, Figure 4. The distance to the three closest neighbours of each RPE nucleus was calculated using the Imaris software. The Spot procedure allowed allocating manually a spot positioned at each RPE nuclei using Otx2 labelling. The position of the spots was adjusted as necessary to align with the centre of each nucleus. The ‘Spots to Spots Closest Distance’ module was used to calculate the average distance to the three closest neighbours for each nucleus within the subset of spheres.

#### Image stack processing for cell tracking

Related to Figure 3I. Hyperstacks used for tracking analyses were derived from a previously published dataset in which nuclei were labelled with H2B-mCherry (Devos et al., 2021). Images were acquired in 8-bit format with a voxel size of 0.3 µm in x-y and 1 µm in z, and a temporal resolution of 2 min 30 s per frame. For the present analysis, image crops encompassing OC morphogenesis from 18.5 to 25.5 hpf were generated. To improve image quality and facilitate tracking, several pre-processing steps were applied: pixel intensities within each stack were homogenized using contrast enhancement (0.35% saturation), and 3D drift correction was performed to improve spatial alignment over time. Processed hyperstacks were exported in H5 format for subsequent analysis using the Mastodon plugin in Fiji (Girstmair et al. 2025).

#### Cell tracking and internuclear distance quantification

Related to Figure 3IJ. To quantify dynamic changes in internuclear spacing during OC morphogenesis, we tracked RPE nuclei in the right and left optic vesicles of two embryos per morph using the Fiji plugin Mastodon. This plugin enables visualization of hyperstacks in multiple orthogonal views, facilitating reliable identification of individual nuclei across successive time points and accurate placement of spots. RPE nuclei were identified based on their position and characteristic morphology. For distance measurements, we tracked neighbouring RPE nuclei to derive internuclear distance vectors. To ensure accuracy, we manually curated and verified the tracks included in the trajectory analyses twice. For internuclear distance quantification, three-dimensional (x, y, z) coordinates of nuclear spots were automatically extracted from Mastodon. Distances between nuclei over time were calculated in 3D space using the Euclidean distance formula.

#### RPE cell surface area

Related to Figure 5. The proximal RPE cell surface areas were obtained by manually tracing the cell contours using the ZO-1 signal as a template in SF. This was done in 3D stacks on OC that were slightly younger than 24 hpf. In these samples, the ventricle space appeared larger, reducing the distance between the two facing epithelial apical sides and making it possible to discern RPE cell outlines. Because of the curvature of OCs, manual tracing was performed on several consecutive slices using depth-colour coded maximum intensity projections to capture the RPE border as perceived by the eye. Although this approach introduced a minor bias, since the 2D projections did not fully account for tissue curvature, we considered it negligible given the large size difference observed between proximal and distal cells.

For the distal RPE cells, the eye could perceive the overall shape, but it was not possible to trace the exact contours rigorously. As only a limited number of cells were available for segmentation, we developed a virtual method to evaluate cell size and shape using Voronoi surfaces. These surfaces were obtained using the Voronoi Fiji plug-in, which automatically generated the surface areas from the nuclei centroids. These latter were automatically calculated from the corresponding 2D segmented nuclei (see below). We also generated Voronoi surfaces for proximal cells (Figure 7).

#### RPE nuclear segmentation (2D)

Related to Figure 4-6. RPE nuclei were segmented manually in Fiji using the Otx2 signal from maximum intensity projections of the distal and proximal regions. The ZO-1 signal served to distinguish RPE from RNE nuclei in the distal region. The region of interest (ROI) corresponding to each nucleus was outlined using the pencil tool (width 1 px). The resulting ROIs were saved as masks for subsequent analyses (see below).Surface areas were calculated automatically from each nuclei ROI using the *Area* plugin in FIJI.

#### RPE volumetric nuclear size (3D)

Related to Figure 6. The Surface module in Imaris was used to measure nuclear RPE volumes. Due to the thickness of the OC, fluorescence signals from the proximal region are attenuated when embryos are imaged in distal orientation. Because segmentation algorithms depend on signal intensity thresholds, this result in under-segmentation and inaccurate volume estimation. To control for this effect, nuclei were analysed from separate samples mounted in distal or proximal orientations, thereby ensuring comparable signal intensity and consistent segmentation of Otx2-positive nuclei. For each sample, we created a Surface mask using the default mode (settings: volume display). The segmentation process, applied to the Otx2 channel, involved a smoothing grain of 0.541 µm for estimated object diameter, thresholding (absolute intensity) and the separation of touching surfaces (split). During the procedure of segmentation, surfaces corresponding to RNE nuclei were also created. These corresponded to uppermost distal neuro-epithelial nuclei (at 24hpf) or proximal neuro-epithelial nuclei (CF 30hpf), which both also accumulated Otx2. These surfaces were discarded manually from the masks. See also Nuclear DAPI intensity quantification.

#### Nuclear aspect ratio and circularity

Related to Figure 5. To quantify nuclear aspect ratio and circularity (see, for example, Figures 5A and 5E), we used the *Shape Descriptors* plugin in Fiji on 2D objects corresponding to segmented nuclei (see above). The regions of interest (ROIs) for each nucleus were derived from the segmentation masks obtained from maximum intensity projections. The aspect ratio was automatically calculated as the ratio of the major to minor axis lengths, while circularity was computed as 4π×area/perimeter^2^. A value of 1.0 indicates a perfect circle, and lower values correspond to increasingly elongated shapes.

#### Nuclear orientation (deviation angle)

Related to Figure 5. Nuclear orientation was quantified as the angular deviation between the nuclear principal axis and the closest OC meridian. Analyses were performed on 2D maximum intensity projections of distal and proximal regions. First, a custom reference pattern consisting of six concentric circles and 48 evenly spaced meridians was generated. For each sample, the reference pattern was pasted into a mask channel and centered on the lens based on the DAPI maximum intensity projection. Segmented RPE nuclei outlines were pasted into a separate channel. For the distal region, only nuclei belonging to the first four ‘rows’ of the RPE were retained for analysis. For each nucleus, the principal axis was obtained in Fiji using the Fit Ellipse function, which provides the orientation angle of the major axis relative to the x-axis (0-180°). The angle of the nearest meridian was determined by drawing a meridian line parallel into the object (nuclei) and retrieving its angle buy manual measure. Given that the Fiji plugin measures orientation angles within a 0-180° range, we corrected the values for nuclei located below the dorsoventral central horizontal line (not shown), which should instead fall between 0 and −180°. In addition, because some nuclei—particularly in the proximal region—displayed a nearly circular shape, only nuclei with an aspect ratio (AR) greater than 1.2 were included in the statistical analyses. The deviation angle α for each nucleus was then calculated as the absolute difference between these two angles (|α-nucleus − α-meridian|).

#### Nuclear DAPI intensity quantification

Related to Figure 6. To test the endoreplication hypothesis, proximal RPE and adjacent RNE nuclei were manually segmented (Figure 6E’). Region-specific substacks were generated from the original z-stacks, and a dedicated mask channel was created in addition to the DAPI and Otx2 channels. Nuclear boundaries were manually delineated in this mask channel on each optical section where DAPI and/or Otx2 signal was detected, enabling 3D reconstruction and quantitative analysis. In this analysis, nuclear volumes were expressed relative to adjacent RNE nuclei rather than as absolute values, in order to control for variability in staining intensity between samples. DNA content was assessed as a proxy for potential endoreplication by measuring the integrated DAPI fluorescence intensity of individual nuclei from both cell types in the same region. For each nucleus, nuclear area, mean DAPI intensity and integrated DAPI density were measured for each optical section. Values were then summed across all sections to obtain the total nuclear volume and total DAPI fluorescence signal intensity.

#### OC size measurement

Related to Figure 7. OC volume was quantified in FIJI using the Measure Stack plugin. ROI outlining the OC were manually drawn on selected key slices of the z-stack. The plugin interpolated ROIs linearly between key slices to reconstruct the object across consecutive sections. The summed measurements across all interpolated ROIs were used to calculate the total volume of the OC.

#### RPE cell number

We obtained the entire RPE structure from a small number of samples. Spots were created manually in Imaris for each nucleus throughout the tissue, using the Otx2 channel and 3D and slice views.

### Measurements for statistical analyses

#### Distance from ZO-1 Margin to lens

Related to Figure 1: For each sample, 15-20 measurements of the d parameter were taken to compute the mean distance.

#### Width of the ventricles

Related to Fig. S1. The average distance between the two ZO-1 apical surfaces was calculated for each morph by taking measurements at 40-70 on each sample.

#### Nearest neighbour distances (NND3)

Related to Figure 3D: For each sample, 60-70 nuclei were analysed in both the distal and proximal regions to calculate the average NND3 for each morph.

Related to Figure 3G: For the three regions analysed, 30-60 nuclei (SF, 24 hpf) and 15-40 nuclei (CF, 24 and 30 hpf) were used per sample to determine the average NND3.

Related to Figure 3H: For each sample, 30-45 nuclei were analysed in the two distal regions to calculate the average NND3 in SF.

Related to Figure 4C: For each sample, 110-160 nuclei were analysed in the proximal region to calculate the average NND3 for each morph.

Related to Figure 4K: For each sample, 50-70 nuclei were analysed in the proximal region to calculate the average NND3 in SF.

#### Internuclear distance (live imaging)

Related to Figure 3J: For each optical vesicle, 10 to 15 pairs of nuclei were analysed in order to calculate the average internuclear distance in both morphs (with the exception of the right optic vesicle of one of the two CF embryos, for which 4 pairs of nuclei were analysed). We have plotted the average internuclear distance per optic vesicle.

#### Nuclear shape and orientation

Related to Figures 5IJ and M-O: The average aspect ratio and circularity of each sample were calculated using 70-90 nuclei in the distal region and 35-50 in the proximal region. To calculate the average deviation angle in the proximal region, we used 25–35 nuclei with an aspect ratio >1.2.

#### Apical cell surface areas

Related to Figure 5D: The average surface area of each sample was calculated using 24-67 Voronoi surface areas from the distal region and 7-18 segmented 2D surfaces from the proximal region.

#### Nuclear size in 2D

Related to Figure 4F and 5H: For each sample, the average surface area of the nuclei was calculated using 20-30 segmented nuclei from the distal and proximal regions.

#### Nuclear size in 3D

Related to Figure 6C: For each sample, the average volume of the nuclei was calculated using 40-60 segmented nuclei (Otx2 channel) from the distal and proximal regions.

Related to Figure 6DF: The average volume of the nuclei was calculated for each sample using six to eight manually segmented RPE and RNE nuclei (DAPI channel) in the proximal region.

## Statistical tests

Given the experimental constraints we aimed to obtain a sample size large enough (N ≥ 5 OCs, indicated in the bottom of each graph) to allow testing statistical significance by using a Mann-Whitney test (unequal variance, *P ≤ 0.05, **P ≤ 0.001, ***P ≤ 0.0001). The number of samples and P-values are either indicated in the figures or the respective legends. For each experiment, numbers in parenthesis indicate biological replicates, meaning the number of biological specimens evaluated (the number of OC). For Figure 3J, we used the Two-Ways ANOVA test.

## Data representation

The data for each measured parameter were exported to an Excel file. Data were plotted in Graph Pad (Prism 9). In box plots, the median is indicated by a central thick line while the interquartile range (containing 50% of the data points) is outlined by a box. Whiskers indicate the minimum and maximum data range; outliers are indicated by a black rhomb and were excluded from further processing.

## Acknowledgments

We thank Stéphane Père for taking care of our *Astyanax* colony. We are also grateful to the Image Analysis Unit at the Pasteur Institute. We would like to thank Jean-Yves Tinevez for his advice on using the Mastodon plugin and processing hyperstack images and Marvin Albert for his guidance on quantifying nuclear orientation. We thank François Graner for his advice on quantifying deviation angles. We would also like to thank Pooja Pullarkat for her help with acquiring the data for Figure 6HI. Finally, we would like to warmly thank Bruno Monier for his critical review of the manuscript.

## Author contributions

F.A. designed the study. M.P., D.V., P.Z., E.G. and F.A. performed the experiments. F.A. performed the quantifications, analyses, and interpreted the data. F.A. built the figures with contributions from J.T-P. and S.R. J.T-P. built the schematics (Fig. 1 and Fig. 8). F.A. wrote the manuscript with contributions from S.R. and J.T-P.

## Movies

Movie 1: Confocal image stack animation showing ZO-1 staining in the entire optic cup in SF (left) and CF (right) at 24 hpf. The animation goes from distal to proximal. ZO-1 accumulates at the apical side of both the RPE and the RNE, thus counterstaining the ventricle. In the CF, the borders of some evl cells (enveloping layer) are seen.

Movie 2: 3D animated rendition of the stack shown in Movie 1, allowing comparison of the shape of the ventricle (and global optic cup shape). The difference in size between the two morphs at the same stage is evident. In SF, optic cup folding is more advance than in CF.

Movie 3: Confocal image stack animation showing the merged staining of ZO-1 (green), Otx2 (magenta) and DAPI (grey) in the entire OC in SF at 24 hpf. Otx2 staining reveals differences in the organisation and shape of RPE nuclei along the proximo-distal axis. DAPI staining reveals the high nuclear density in the RNE, as well as the proximity of the outermost nuclei of the pseudostratified RNE and the RPE.

Movie 4a and 4b: 3D animated renditions comparing the anatomy of the RPE and optic cup at 24 hpf in the two morphs. Otx2 staining reveals regional differences in the organisation and shape of RPE nuclei. ZO-1 staining reveals the 3D shape of the RPE-RNE apical interface and the optic stalk. This marker also reveals the polygonal shape of the enveloping layer’s cells. Of note, normalization (0.35%) was applied to the entire ZO-1 image stack channel to increase the signal in the proximal region and highlight the 3D shape in SF.

Movie 5: 3D animation illustrating the organisation and density of RPE nuclei across an entire optic cup in SF at 24 hpf. Colour-coded (blue to magenta) 3D map of NND3 across the RPE surface reveals the graded patterns of nuclear density along the different axes. Note the differential density of RPE nuclei along the proximo-distal axis. Note as well the differential spot density in the ventral and dorsal domains of the distal region (turquoise spots).

Movie 6: 3D animation showing the three zones used to quantify the density of RPE nuclei along the proximal-distal axis in an SF at 24 hpf. The Otx2 channel was used to create grey spots at each RPE nucleus position. The colour code distinguishes between the distal (yellow), intermediate (red), and proximal (green) regions of the RPE. Note the reduced spacing of the yellow spots in the distal region and the increased spacing of the green spots in the proximal region. A gradual spacing of the spots is manifest along the proximal-distal axis.

Movie 7: 3D animation illustrating the organization and density of RPE nuclei across an entire optic cup in CF at 36 hpf. Colour-coded (blue to magenta) 3D map of NND3 across the RPE surface reveals the graded patterns of nuclear density along the different axes. Note the extended nuclear density of RPE in the ventral region compared to 24 and 30 hpf in CF.

Movie 8: A 3D animated volumetric rendering showing nuclear organisation in the proximal region of the SF optic cup at 24 hpf. The RPE nuclei (labelled with Otx2 in magenta, some of which are outlined in yellow) are circular in shape. Just below, the RNE nuclei stained with DAPI (some of which are outlined in turquoise) can be seen. Note the proximity of the RNE nuclei to the RPE nuclei. Also observe the large Otx2-negative nuclei. These are likely to correspond to neural crest cells. The volume of tissue shown corresponds to a depth of 18.6 µm.

---

**Fig. S1:**
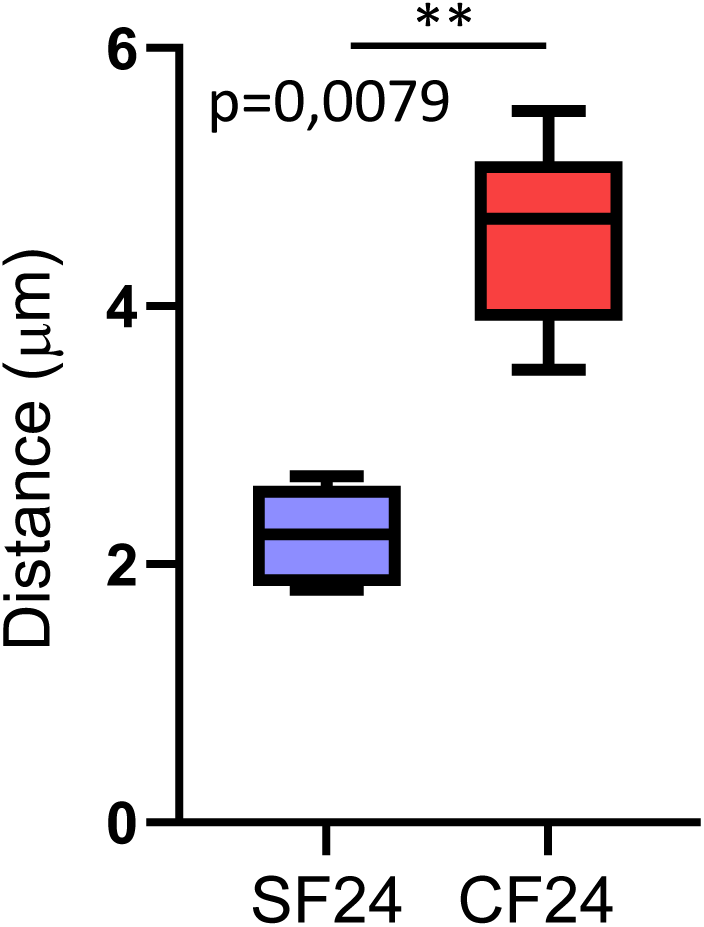
Increased width of the ventricle in the cavefish RPE. Quantification shows the width of the ventricular space that separating the RPE and the retinal neuroepithelium for the two morphs at stage 24.

**Fig. S2:**
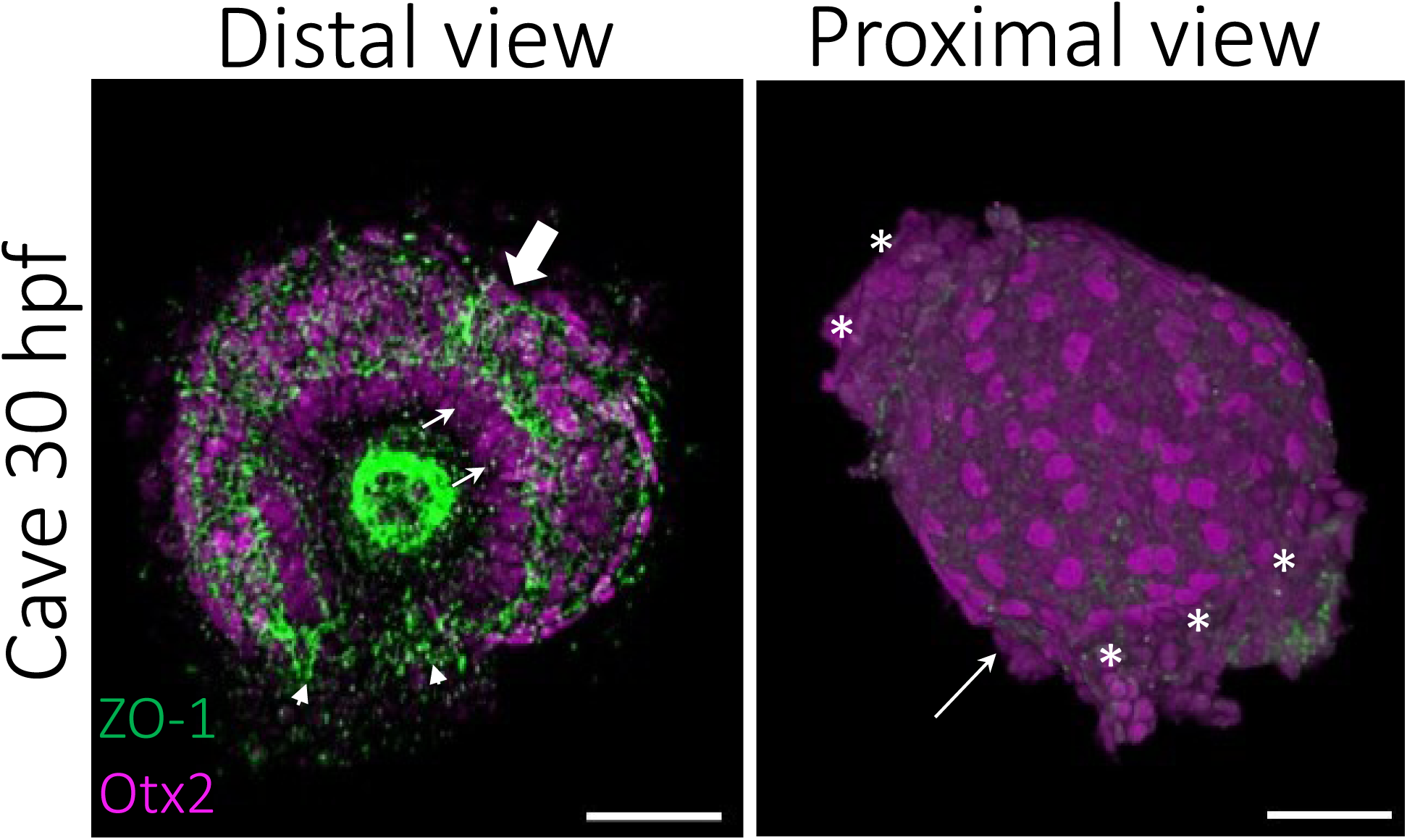
Anatomy of the RPE in CF at 30 hpf. 3D distal and proximal views of an optic cup immunostained for ZO-1 (green) and Otx2 (magenta), showing the regional nuclear organization of the RPE. The orientation of the optic cup is as in Figure 2A. (Left) The arrowheads show where the most ventral ends of the ventricle are located, at the level of the optic fissure. The large arrow shows a superficial fold of the dorsal optic cup, which is visible at this stage. The position of the lens is shown by the strong green signal in the centre. Arrows point to ciliary margin progenitors. (Right). The arrow indicates the position of the optic stalk, which is not shown in the picture. Asterisks indicate tissue outside the optic cup. Observe the irregular arrangement and shapes of the nuclei in the proximal region. Scale bar: 50 µm.

**Fig. S3:**
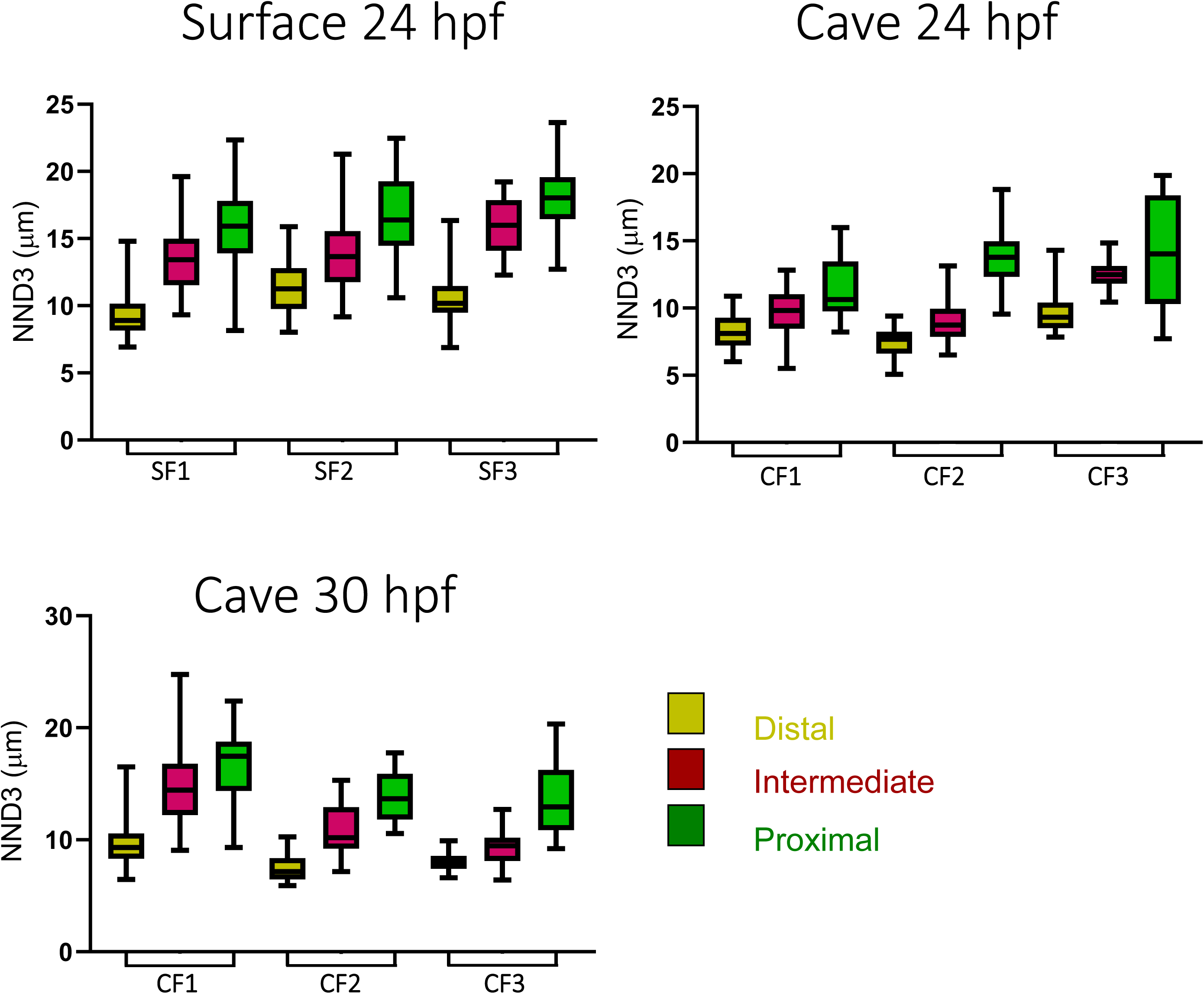
Quantification of NND3 across different RPE sub-regions in the two morphs at indicated stages, in different samples. The colour-code is indicated. Three samples are shown in each graph, showing the reproducibility of the observation and the interindividual variability.

**Fig. S4:**
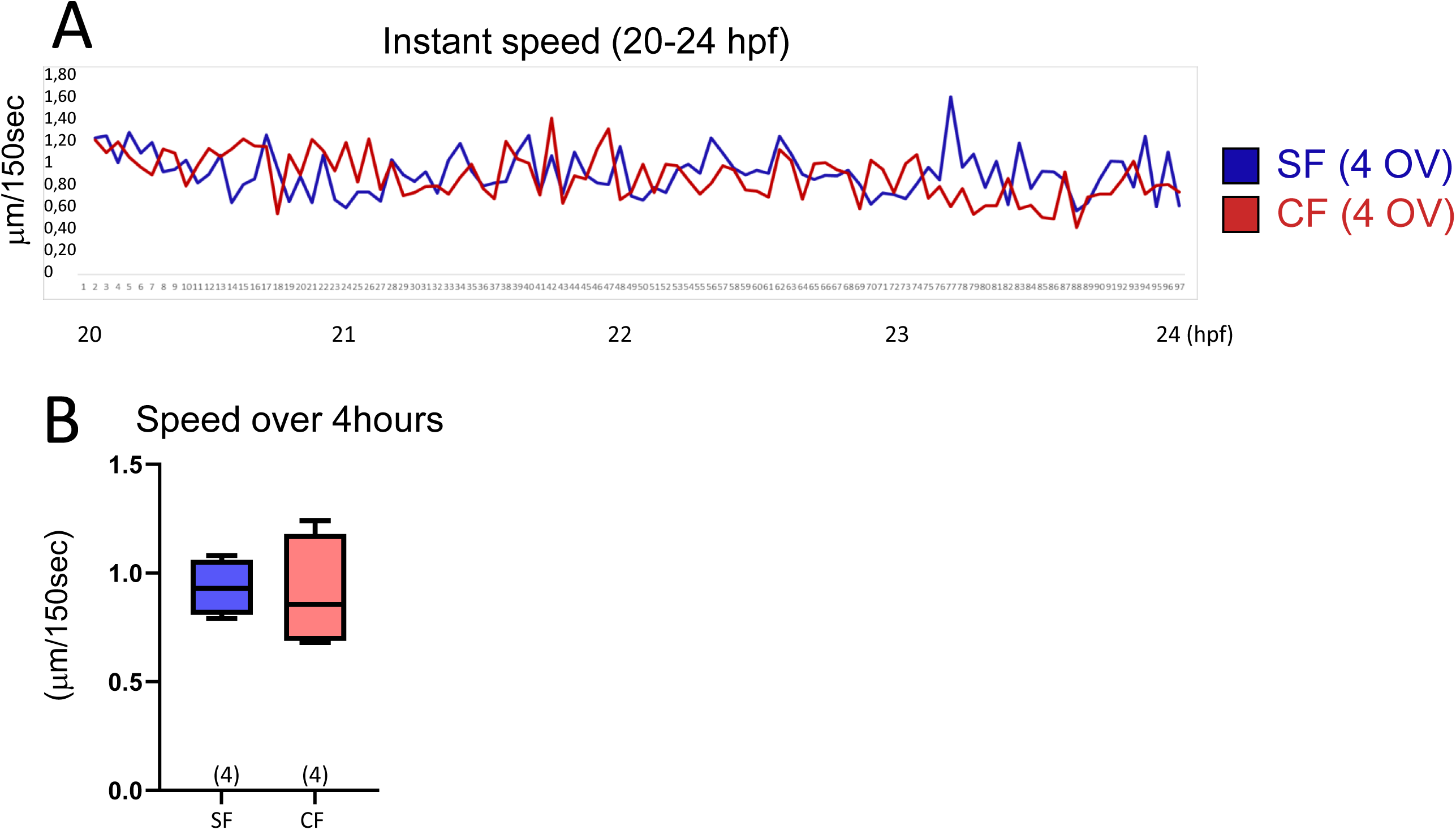
Instant speed and mean speed of RPE nuclei are equivalent in the two morphs. (A) The graph shows the instant speed for the two morphs, with the number of optic vesicle (OV) used for the quantification. Six to eight cells were used for each OV. (B) Boxplot showing the mean of the speed per sample in the two morphs.

**Fig. S5:**
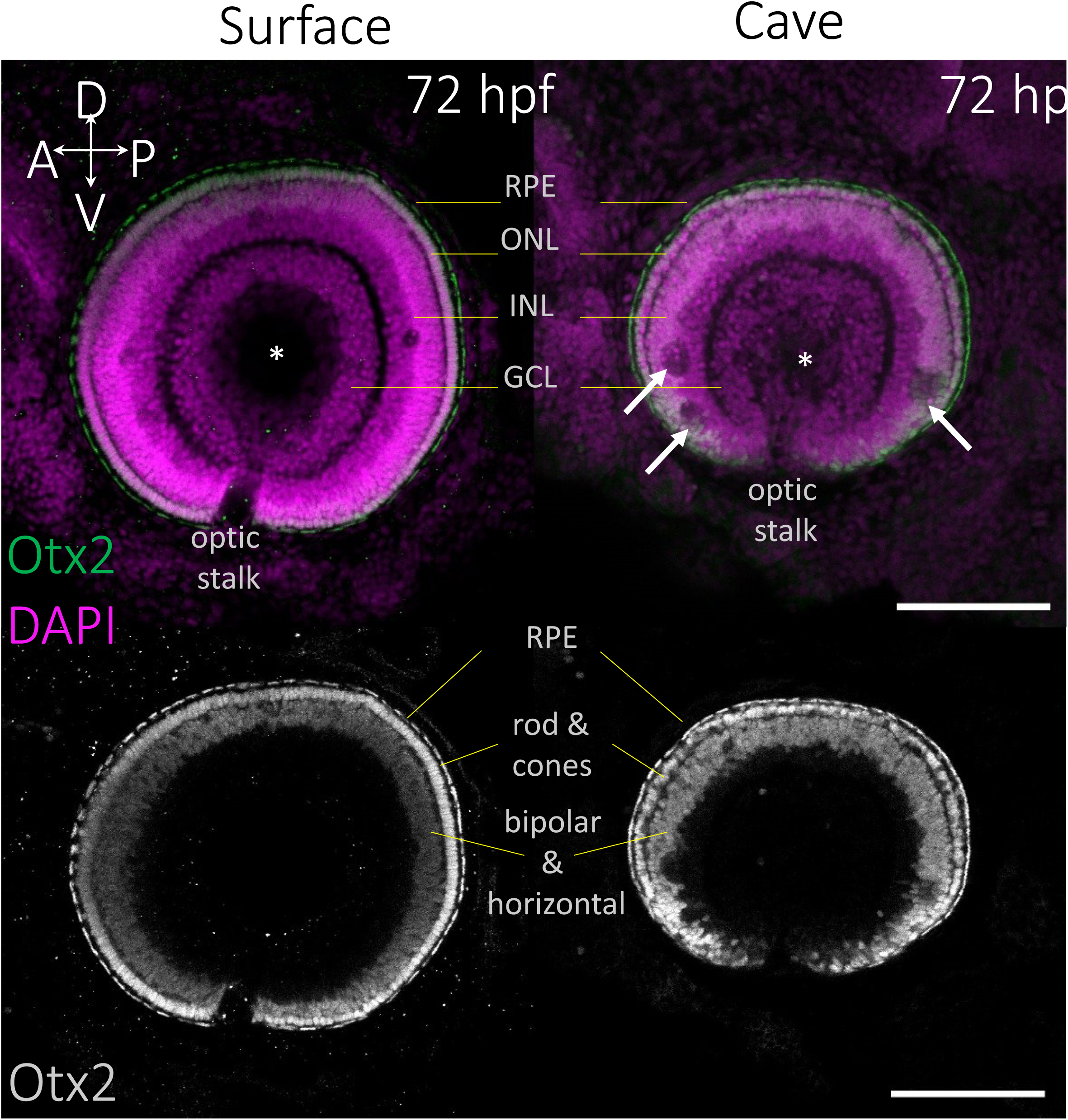
Organisation of the eye at 72 hpf in the two morphs as seen with Otx2 and DAPI staining. All panels show maximum intensity projections of confocal image stacks in the intermediate zone of the eye. At 72 hpf Otx2 is detected in the RPE, the outer nuclear layer (ONL) and the superior layer of the inner nuclear layer (INL). The retinal subtypes corresponding to these layers are indicated in the lower panel. The asterisk indicates the position of the lens. Large arrows indicate clusters of degenerating retinal cells in the CF. Observe the irregular shape and width of the ONL in CF. Also note the reduced size and abnormal ventral shape of the eye in CF.

**Fig. S6:**
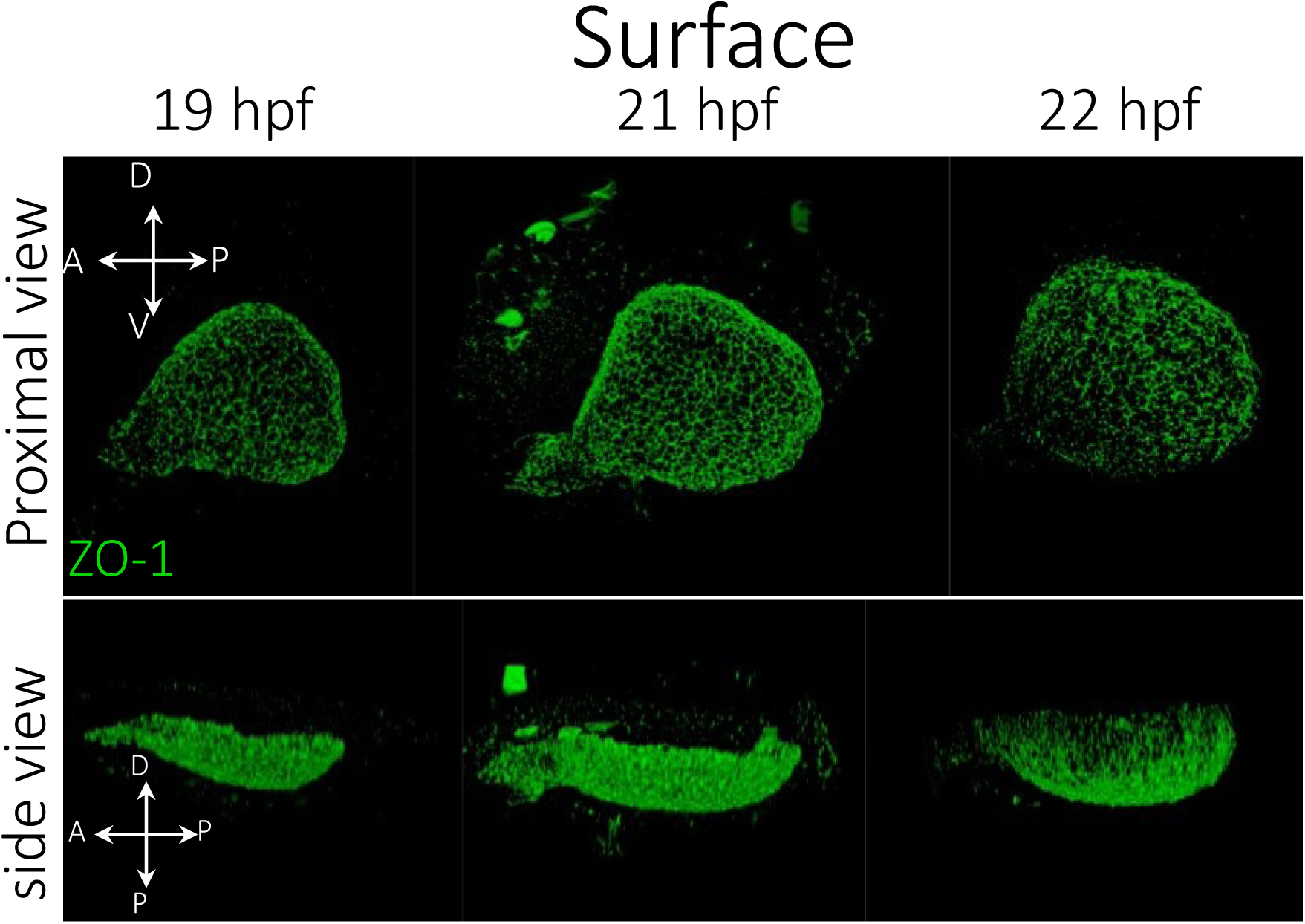
3D renditions of optic cup stained with ZO-1 highlighting the shape change in the OC in surface fish. As illustrated by the proximal view, the optic cup undergoes a progressive rounding process. In addition, the side view illustrates a curving or bending motion in the proximal region. Orientations are indicated. Bottom: D = distal and P = proximal.

**Fig. S7:**
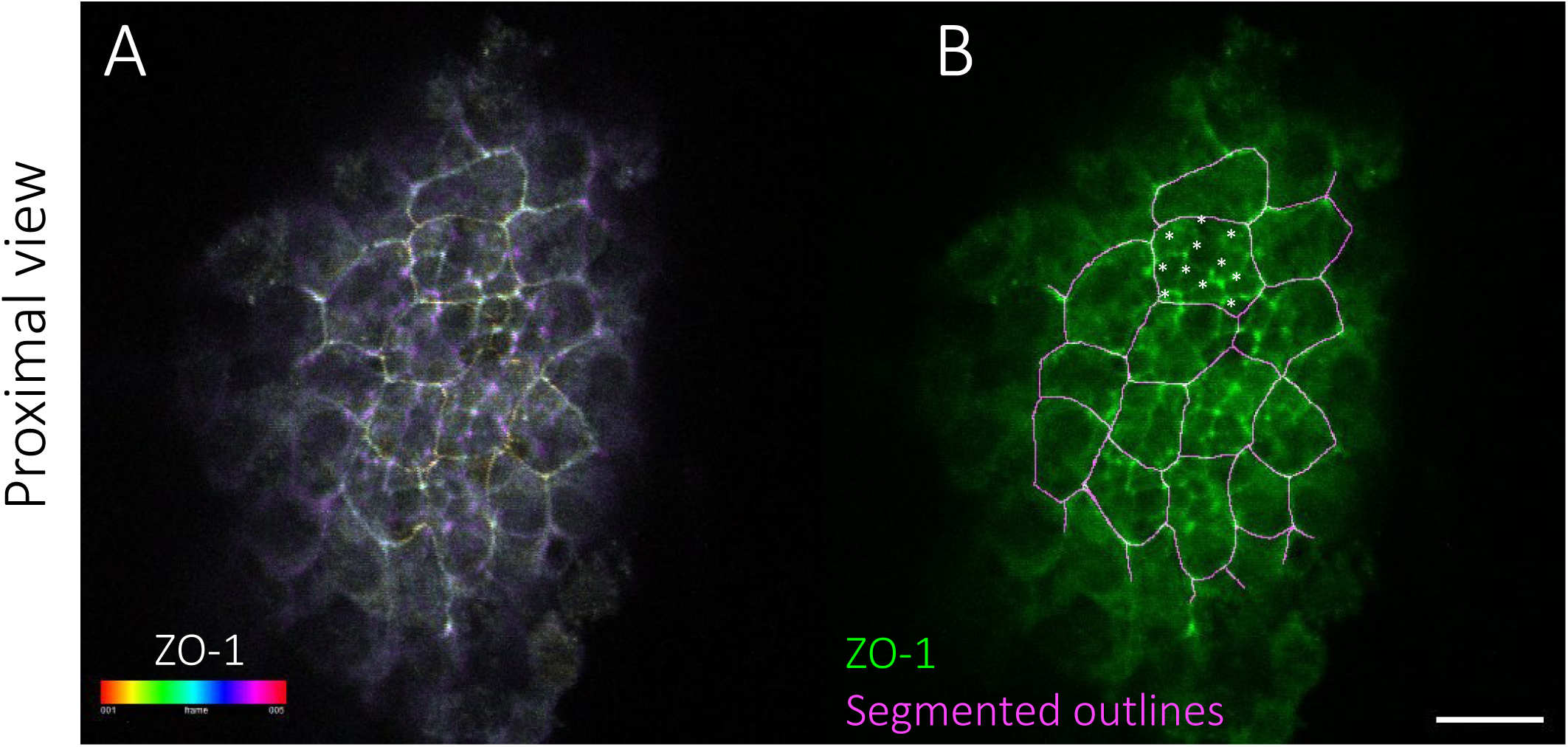
Confocal images showing an overlap of the apical ZO-1 signals from the RPE and the NE in the proximal region. (A) Depth colour-coded maximum intensity projections of 5 optical slices (z-step = 0.42 µm), showing ZO-1 accumulation in the RPE (large cells) and retinal neuroepithelium (below). (B) ZO-1 signal (green) overlaid with manually segmented RPE cell contours (magenta). The apical surface areas of 11 retinal progenitor cells in contact with 1 RPE cell are indicated with asterisks. These images highlight the strategy and method used to identify unambiguously the two cell types, and quantify their size and organization features. Scale bar: 20 µm.

**Fig. S8:**
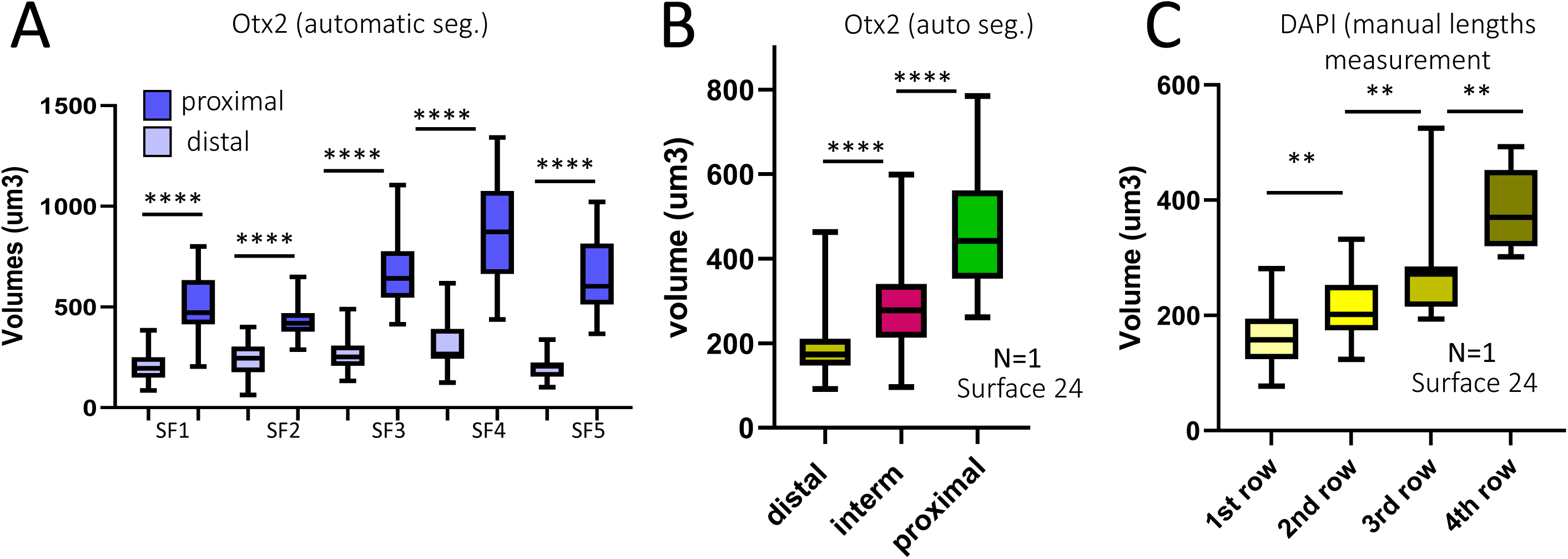
Quantification of nuclear size in various regions of the RPE in SF at 24 hpf using different approaches. (A) Comparison of nuclear volumes in the distal and proximal regions in five samples. The Otx2 signal was segmented automatically using the Imaris software. A significant difference between the two regions was observed for each sample, thereby showing the robustness of the result. (B) Comparison of nuclear volumes between the distal intermediate and proximal region in one sample using the same method as in A. (C) Comparison of nuclear volumes within the four more distal ‘rows’ in one representative sample. Width, height and length in x,y and z were measured for a series of nuclei and the volume was calculated using the formula for an ellipsoid. (D) Comparison of nuclear volumes between retinal neuroepithelium and RPE in the proximal region of the optic cup. The Dapi signal was used to manually segment the nuclei outline using the Fiji software.

**Fig. S9:**
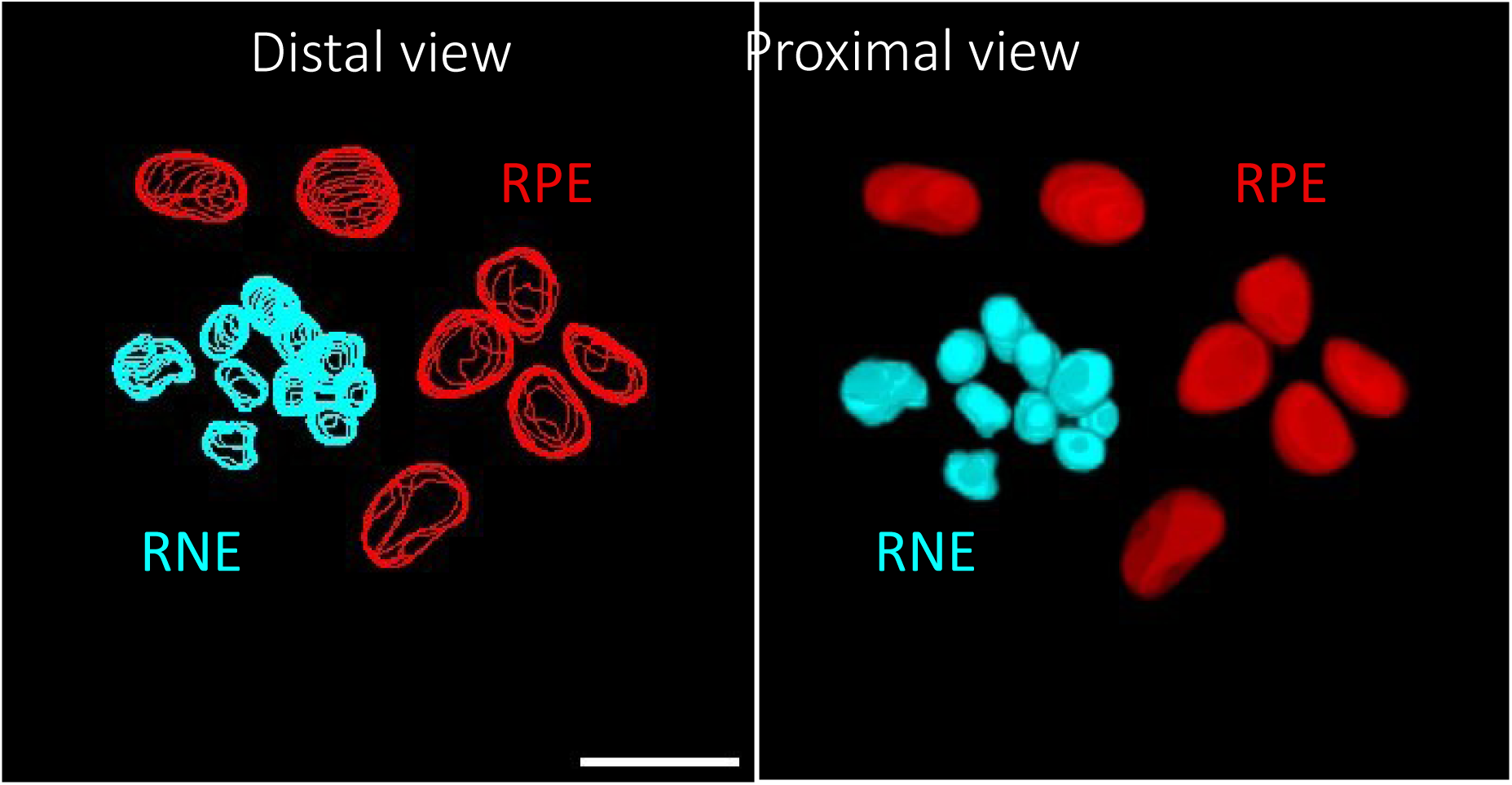
3D views of manually segmented proximal RPE and RNE nuclei in SF at 24 hpf. (Left) Nuclei were manually segmented by tracing outlines in each z plane of the confocal stack. The image shows the maximum intensity projection. (Right) 3D rendering using the 3D viewer plugin (Fiji) from the image stack after the outlines were filled in. A significant difference in size and 3D shape is observed between the two nuclei subtype. The 7 RPE nuclei (red) seen from top display lenticular shapes. The 10 RNE nuclei (turquoise) are much smaller and have elongated shapes. The tissue volume shown corresponds to a depth of 18,3 um. Scale bar 20 um.

